# Brain tissue mechanics is governed by microscale relations of the tissue constituents

**DOI:** 10.1101/2022.10.19.512076

**Authors:** P. Sáez, C. Borau, N. Antonovaite, K. Franze

## Abstract

Local mechanical tissue properties are a critical regulator of cell function in the central nervous system (CNS) during development and disorder. However, we still don’t fully understand how the mechanical properties of individual tissue constituents, such as cell nuclei or myelin, determine tissue mechanics. Here we developed a model predicting local tissue mechanics, which induces non-affine deformations of the tissue components. Using the mouse hippocampus and cerebellum as model systems, we show that considering individual tissue components alone, as identified by immunohistochemistry, is not sufficient to reproduce the local mechanical properties of CNS tissue. Our results suggest that brain tissue shows a universal response to applied forces that depends not only on the amount and stiffness of the individual tissue constituents but also on the way how they assemble. Our model may unify current incongruences between the mechanics of soft biological tissues and the underlying constituents and facilitate the design of better biomedical materials and engineered tissues. To this end, we provide a freely-available platform to predict local tissue elasticity upon providing immunohistochemistry images and stiffness values for the constituents of the tissue.

---

Many biological processes in the brain involve mechanical interactions of cells with their surrounding tissue [1]. Cells exert forces on their environment and probe and respond to its local mechanical properties. Accordingly, CNS tissue mechanics is an important regulator of cell function during development [2, 3] and disease [4]. During normal ageing and pathological processes, CNS tissue constituents change, including the extracellular matrix (ECM), myelin sheets around neuronal axons, and the number of cells [7, 6, 8]. These structural alterations are accompanied by changes in tissue mechanics [5, 6].

During the last decades, extensive experimental tests have been performed to determine mechanical properties of the brain (see, among many others, [9, 10, 11, 12, 13, 14, 15, 16, 17, 18, 19]). Nowadays, there is a clear consensus that the brain exhibits time-dependent behavior [14, 19], both at small and large strains, and that elastic behavior dominates viscoelasticity at the cell and tissue level[20]. The elastic modulus of CNS tissue, a measure of its elastic stiffness, ranges from, approximately, 0.1 to 2 kPa [11, 19, 13]. This range of stiffness has been observed across species and different regions of the CNS [16, 17, 18]. However, while our knowledge of how brain tissue behaves mechanically at the tissue scale has progressed substantially, how individual brain components contribute to macroscopic brain tissue mechanics is still poorly understood.

Most current mechanical models of brain tissue (see [21, 19]) homogenize the behavior of the tissue constituents into a single constitutive law without further consideration of what components are present or how each of them is organized within specific brain regions [9, 10, 11, 19, 13]. Although there has been a large amount of research devoted to understanding the brain’s composition and architecture, this information has so far only sparsely found its way into advanced mechanical models of the brain [22, 23, 24]. However, like in other biological tissues composed of networks of cells and ECM, the arrangement and stiffness of the underlying brain constituents should determine the mechanics of the organ as a whole.

The definition of a Strain Energy Density Function (SEDF) has been the standard approach in the characterization of soft biological tissues [25]. Within this thermodynamic framework, the contribution of each individual tissue constituent is gathered by an additive decomposition of a SEDF per constituent. Naturally, the additive decomposition of the SEDF imposes affine deformations in the tissue constituents. In other words, it implies that the tissue constituents are arranged in parallel to each other. However, such approach may not accurately reproduce how the tissue components actually assemble.

CNS tissue is mainly built of two cell types, neurons and glial cells. Fig. 1(a) presents an overview about the main components of the brain and their distribution in the tissue. Pyramidal and granular neurons represent 90% of all neurons in the hippocampus, and the remaining 10% mainly consist of GABAergic interneurons [26]. Some neuronal axons are wrapped by oligodendrocytes, a type of glial cells. The resulting myelin sheath surrounds axons to improve electrical conductivity, and it is characterized by a comparatively high stiffness [27, 28, 29]. In addition to oligodendrocytes, other types of glial cells are found in CNS tissue, with astrocytes being the most abundant glial cells in mammals.

**Figure 1:**
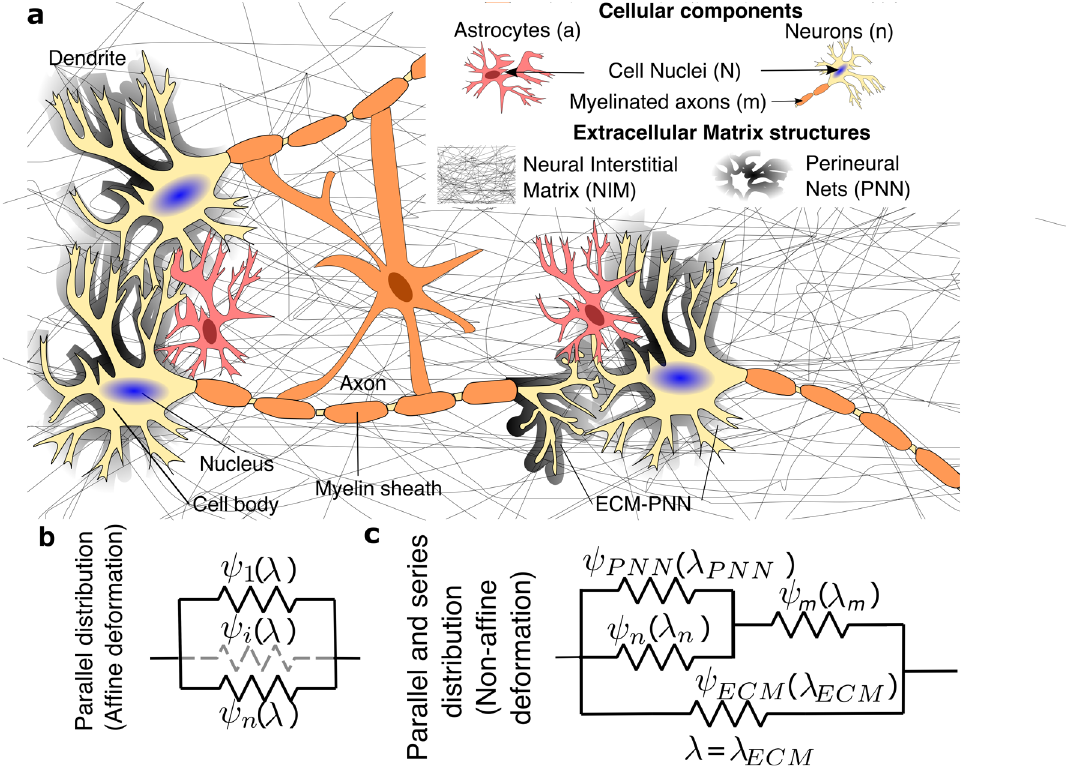
(a) Description of the main brain components. We consider cell bodies, *ρ*_*n*_, perineural nets (PNN, black translucent), *ρ*_*P NN*_, myelinated axons (orange), *ρ*_*m*_, and neural interstitial matrix (NIM), *ρ*_*NIM*_. PNNs surround neuronal cell bodies and dendrites and the NIM surrounds glial and neuronal cells. Neurons, *ρ*_*n*_ are drawn in yellow. Glial cells, *ρ*_*a*_ are drawn in pink. (b-c) Distribution of tissue constituents and modeling of a simplified 1D model based on a classical parallel arrangement (b) and on the distribution presented in this work (c). *Ψ*_*i*_ is the SEDF of each tissue constituent. *λ* and *λ*_*i*_ is the total stretch in the affine scheme (b) and the stretch of each constituent in the non-affine scheme (c), respectively.

The ECM [30] provides adhesion sites for cells to organize into distinct regions. However, unlike in other biological tissues, the ECM of the CNS lacks an abundant collagen network, which is a main determinant of tissue stiffness, and is instead mostly composed of hyaluronic acid (HA) and proteoglycans [31]. Morphologically, the CNS ECM is divided into three main structures that create a scaffold with mechanical and functional roles: perineuronal net (PNNs), the neural interstitial matrix (NIM), and the basal lamina (BL) [32]. The relative ratios of PNNs and NIM components across different brain regions are still not known. In addition to the ECM, blood vessels pass through the tissue.

Recent studies have started to investigate the relationship between the stiffness of specific brain regions and their constituting structures using phenomenological linear regression models [17, 18, 24, 23]. However, although we already know the mechanical properties of individual neurons and glial cells [20, 33, 34, 35], and that the presence of myelinated axons has a clear impact on the mechanics of brain tissue [36, 15, 8], we still do not fully understand how the multiple constituents of CNS tissue ensemble to establish its mechanical behavior at the tissue scale.

Here, we propose a novel mechanical model that explains brain tissue mechanics at different scales. We focus on the hippocampus and cerebellum of the mouse brain,heterogeneous brain regions in terms of stiffness and composition.

## 1 Materials and Methods

### 1.1 Sample preparation, ROI identification, indentation protocol and image acquisition

All experimental protocols and data have been previously described and published (see details in [23, 18] and SI Appendix). For DMA experiments, tips of 60-100um were used to indent the sample up to 10-17um to avoid effects of surface roughness and of sensing individual brain components so that tissue heterogeneities within each ROI were homogenized. The indentation depth was chosen to stay in the small strain regime (*ϵ*_*T*_ *<* 5%). For data analysis, the DMA model was used to fit the cosine function over oscillatory data to extract storage and loss moduli as a function of indentation-depth. The Hertz model was used to fit the initial loading data to obtain the precise the contact point location which is needed for the estimation of contact radius (see SI Appendix).

### 1.2 Image analysis

We analyzed image data using custom-written MATLAB codes which allowed automatic processing of multiple images by providing their corresponding ROIs coordinates. To minimize intensity disparity due to acquisition conditions, we preprocessed all images to adjust their histograms to match the histogram of an arbitrarily chosen reference image. After that, every color channel was individually normalized by its maximum intensity value. Subsequently, for every pixel of each image, the percentage of each color channel intensity (*p*_*r*_, *p*_*g*_, *p*_*b*_) was computed. Then, for each ROI and each channel (c) we calculated the weighted mean 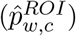 of the color intensity percentage, using the total intensity of each pixel (the sum of the three channel intensities) as weight:

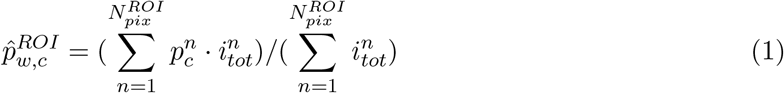

Multiplying the weighted mean intensity of each channel by the total average intensity of the ROI 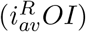 is equivalent to calculating the average channel intensity 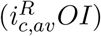, which is later used to weight the storage and loss modulus of each of the brain components (see Supp. Material).

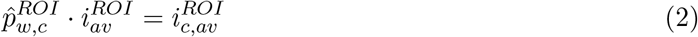

### 1.3 Fitting of material parameters

We used a custom-written MATLAB code to fit the material parameters of the model, i.e E’ and E” for each constituent. We made use of the *fmincon* function to find the minimum of a constrained nonlinear multivariable goal function

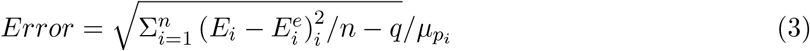

Here *n* is the number of ROIs, *i* represents each ROI and 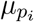 denote the mean of the experimentally measured E’ and E”. In addition *q* is the number of parameters of the model, such that *n* − *q* denotes the number of degrees of freedom.

## Results

### Local hippocampus composition correlates with tissue stiffness

First, we analyzed the local composition of the juvenile mouse hippocampus, using immunohistochemistry. We focused on myelin (using myelin binding protein, MBP, as marker), as an indicator of myelinated axons [8], nuclei of neurons (NeuN), nuclei of all cells (Hoechst), and astrocytes (glial fibrillary acidic protein, GFAP) in different regions of interest (ROIs) of the tissue (Fig. 2a-e and SI Appendix, Fig. S14)), as reported elsewhere [23]. Large variations in densities of the individual markers over the different ROIs where observed [23] (Fig. 2f and Table S1).

**Figure 2:**
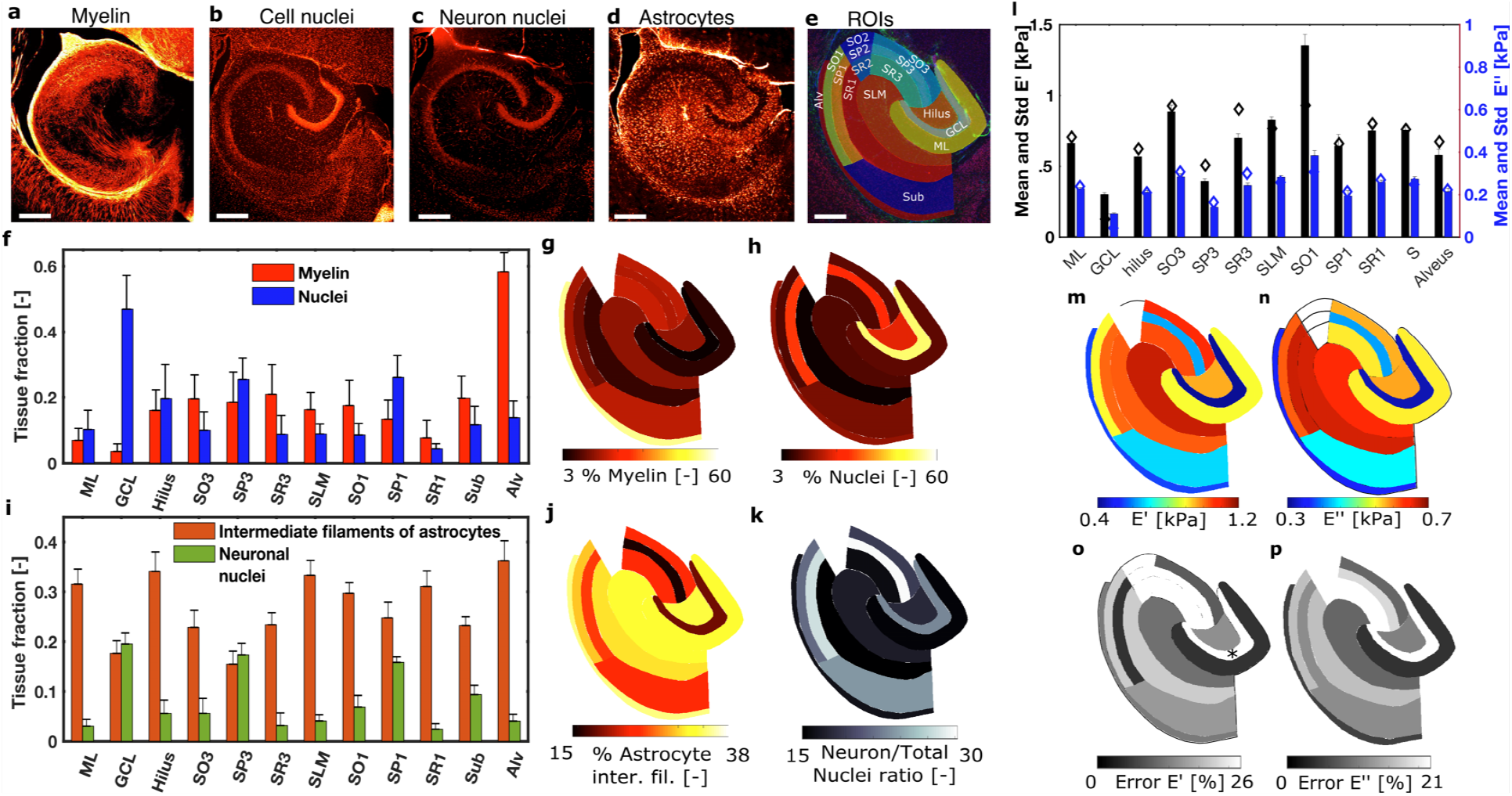
Imaging and quantification of brain tissue composition and stiffness values of juvenile mice. Imaging of (a) myelin (MBP), (b) cell nuclei (Hoechst), (c) nuclei of neurons(NeuN) and (d) intermediate filaments of astrocytes (GFAP) in specific regions of the hippocampus. (e) Abbreviations used for regions are: Alv - alveus, Sub - subiculum, SLM - stratum lacunosum moleculare, SR - stratum radiatum, SP - stratum pyramidale, SO - stratum oriens, ML - molecular layer, GCL - granule cell layer. White regions indicate excluded due to similarity with adjacent regions. Scale bars (a-e) are 200*μm*. Mean and standard deviation of the quantificatied tissue components (f-k). Mean and standard deviation of the myelin and nuclei (f) and representation of the mean value in each specific ROI (g-h). Mean and standard deviation of the neuron nuclei and intermediate filament of filaments of astrocytes (i) and representation of the mean value of intermedia filaments and the ratio between neuron vs. total nuclei in each specific ROI (j-k). (l) Mean and standard deviation of E’ and E” for the different regions of the hippocampus. Diamond marks shows the prediction of the proposed mechanical model and bars show the measured values. (m-n) Representation of E’ and E” in all the ROIs. (o-p) Error of the proposed mechanical model for E’ and E” represented in each ROI.

The granular cell layer (GCL), which mainly consists of neuronal cell bodies, reached the highest concentration of nuclei with ∼46%, while the amount of myelin was just a residual 0.5%, the lowest of all ROIs (see Materials and Methods). The Alveus (Alv), on the other hand, which is a region containing mainly myelinated fibers that cover the ventricular parts of the hippocampus, had the highest concentration of myelin, ∼50%, while the amount of cell nuclei reached 15%. All other regions presented a more uniform distribution of tissue components, in agreement with previous image analysis results [23] (for details see Fig. 2f-k and Table S1).

In order to quantify the fraction of neuronal and glial cell nuclei, the distributions of NeuN and the astrocyte marker GFAP were measured [18, 23]. ROIs with higher amounts of cell nuclei (GCL, Sub and SP3, Fig. 2f) had the largest ratio of neuronal nuclei to total number of nuclei (see Fig. 2k) and showed a low concentration of GFAP (Fig. 2 and Table S1). Other regions, including the ML, SR1, SR3, SLM, Hilus and Alveus, were characterized by a low ratio of neuronal/total cell numbers and a high concentration of GFAP, confirming previous descriptions of the spatial heterogeneity of brain tissue components in the hippocampus [7].

To account for the remaining, non-cellular components of the tissue, we made the following assumption for the fraction of ECM and, in particular, of PNNs and NIM: *ρ*_*ECM*_ = 1 −(*ρ*_*N*_ + *ρ*_*m*_), that is, the space not occupied by either nuclei (*ρ*_*N*_) or myelin (*ρ*_*m*_) is assumed to be ECM. Then, we assume that NIM is more prevalent in regions where more astrocytes (*ρ*_*a*_) are present and PNNs where more neuronal nuclei (*ρ*_*n*_) are localized. The mechanical contribution of the vasculature is included within the NIM. Therefore, we defined *ρ*_*NIM*_ = *ρ*_*ECM*_ * *ρ*_*a*_/(*ρ*_*n*_ + *ρ*_*a*_) and *ρ*_*PNN*_ = *ρ*_*ECM*_ * *ρ*_*n*_/(*ρ*_*n*_ + *ρ*_*a*_) (SI Appendix, Fig. S1-3).

Then, we compared recently published data of the stiffness of each ROI (see Materials and Methods and [23] for further details) with the distribution of each ROI’s components. Tissue stiffness had been measured using a nanoindentation approach, where forces were applied that were similar in magnitude and time scale as the forces exerted by CNS cells [2, 37].. Means and standard deviations of the storage moduli E’, which are a measure of the tissue’s elastic component, for each ROI at every indentation level, *δ*, are provided in Fig. S1. We focused on an applied *δ* = 13*μ*m (Fig. 2(l-m)), but all the features described next were consistent along the different *δ*(SI Appendix, Fig. S10, Tables S12-13).

In summary, the lowest value of E’ ∼0.3kPa was obtained for the GCL. As shown in Fig. 2, the GCL is a region mostly occupied by neuronal cells, specifically granular cells and astrocytes, with a comparatively small amount of myelinated axons (5%). The SP3, which mostly consists of pyramidal neurons and a moderate amount of myelinated axons (20%), was the second softest region (∼0.4kPa). These two regions were characterized by the highest fraction of neuronal nuclei in the hippocampus. Surprisingly, the third softest region was the Alveus (∼0.7kPa), a region containing almost 60% of myelinated axons, few neuronal cell bodies, and a substantial amount of astrocytes. Most other ROIs fell in the 0.8-1.2 kPa range. The stiffest ROIs was the SO1, with E’ ∼ 1.3kPa, which has a moderate fraction of astrocytes (30%) and low amounts of other stained components.

To analyze how the tissue composition regulates tissue rigidity, we first investigated the correlation between the tissue fractions (Fig. 2) and the stiffness of all ROIs (SI Appendix, Fig. S4 and Table S2). We observed negative correlations between structure and stiffness for all tissue constituents except for the fraction of astrocytes, which showed a positive correlation. Only the correlations between tissue stiffness and the densities of all nuclei and those of neurons alone had a significant Pearson’s correlation coefficient (r=-0.69 and r=-0.66, respectively), in agreement with previous data [23]. These results indicated that the characteristic mechanical signature of each ROI may be related to the local tissue composition, and that tissue mechanics likely results from a combination of all the constituents rather than being dominated by one single component.

### Brain mechanics cannot be explained through affine deformation of its components

To test this hypothesis, we set out to define a constitutive law describing the stiffness of each ROI by incorporating information about the local tissue composition. Previous work correlating stiffness and tissue composition in the nervous system developed phenomenological models assuming linear and non-linear relationships to reproduce composition-stiffness relations [22, 23] or assuming that only the tissue component with the highest correlation with the tissue stiffness contributes to its rigidity [24]. Here, we derived a homogenized model, based on a mean-field (MF) approach, to predict brain tissue mechanics as a function of the distribution and stiffness of each tissue constituent (see Materials and Methods, point 1.1 for details on the representative volume element [39]). We first defined a MF model based on the total SEDF Ψ_*T*_ of the system, the density energy required to deform the tissue under an external load (we provide a complete derivation in the SI Appendix). This is a standard approach in modeling soft biological tissues (e.g. arteries [25, 40], cartilage [41] and brain tissue [21]). Within the small strain limit, and assuming that the tissue is isotropic, almost-incompressible, and that bonds between constituents are strong so that slippage does not occur at small strains [38], we can define the stress-strain relations as a function of the storage Young’s modulus E (a measure of tissue stiffness). The total equivalent *E*_*T*_ of the tissue, that is the total stiffness of the tissue due to the presence of several constituents with Young’s modulus E_*i*_ and fraction *ρ*_*i*_, is 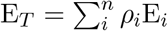, which recovers the classical Voigt estimation [42].

Following the additive decomposition of the SEDF, i.e. the Voigt estimation, we took the values of tissue composition quantification (Fig. 2) and performed a fitting procedure against the experimental values of E’ at different indentation levels (see Materials and Methods). We fixed the storage modulus of cell nuclei to E’_*N*_ = 0.6kPa and took the modulus of myelinated axons, E’_*m*_= 2.0kPa, and of the PNN and NIM, E’_*P NN*_ =1kPa and E’_*NIM*_ =1kPa, respectively, as initial seeds of the fitting procedure, in agreement with previously published values [20, 35, 43, 33, 6]. The fitting procedure yielded mean errors in all regions and indentation levels of ∼21% based on the fraction of their components (SI Appendix Fig. S10 and Tables S12-13). E’_*m*_ increased linearly from ∼0.3kPa at *δ* = 6.5*μ*m up to ∼1.3kPa at *δ* = 17*μ*m, and E’_*P NN*_ and E’_*NIM*_ also increased linearly from ∼0.5kPa and 0.2kPa up to 1.7kPa and 0.7kPa, respectively.

As mentioned above, the additive decomposition of the SEDF imposes a parallel arrangement of the brain constituents and, therefore, an affine deformation of the tissue constituents. In such affine deformations, the stiffest component dominates the mechanical response of the tissue. The stiffest component of the brain, according to literature, are myelinated axons, with elastic moduli between ∼2kPa-1MPa [44, 20]. However, the fitted E’ values for myelin (∼0.2-0.6kPa, SI Appendix, Fig. S12) were lower than these published values. Even if we consider the lowest range of the myelinated axons, stiffness values of 2-20kPa, and impose it into the additive decomposition of the SEDF, it would lead to an equivalent stiffness of the tissue of ~3-12kPa (the component stiffness multiplied by its tissue fraction). This resulting stiffness of the tissue represents errors of ∼300 - 1500%, 1 - 2 orders of magnitude higher than the errors found when the model parameters were freely fitted by the minimization algorithm, suggesting that affine deformations cannot explain the observed behaviour. Moreover, it can be argued that tissue components are not all connected in parallel. Neurons have clearly differentiated regions (e.g. cell bodies and the myelinated axons) with specific mechanical properties [20], which must deform in a non-affine manner under an imposed force.

These findings and arguments suggest that the mechanical response of the brain could not follow an additive decomposition of the SEDF or, equivalently, be explained through affine deformations of its constituents.

### Non-affine deformations of brain components explain tissue mechanics

To evaluate possible explanations of the inconsistencies described above, and to derive a physical model capable of solving them and predicting local tissue stiffness based on the mechanical properties and arrangements of its constituents, we took a closer look at the brain tissue structure (Fig. 1). Some of the tissue constituents do not assemble in the tissue in a parallel arrangement but rather in series, as for example myelinated axons and their somata. Thus, the modeling limitations described for the additive decomposition of the SEDF could arise from the inherent affine deformation imposed on the tissue constituents. If tissue constituents arrange in series, forces are transmitted continuously in all the elements and the constituents will be subjected to non-affine deformations. We can generalize this idea within a consistent thermodynamic formulation, taking advantage of the complementary energy density function (cSEDF), 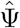 [45, 46]. In terms of the reduced storage modulus, the equivalent *E*_*T*_ of *n* components arranged in series, i.e. defined through its cSEDF, is 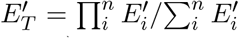, which recovers the Reuss estimate [47] (see SI Appendix for a complete derivation). We particularized this approach to our problem and modeled the structural organization as follows: PNNs are linked in parallel with the neuronal somata, which are linked to the myelinated axons in series. These two elements in series, on the other hand, are linked with the NIM in parallel (see Fig. 2). Therefore,

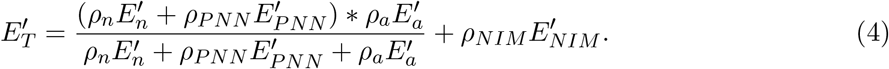

We then used this structural description of the tissue to analyze whether we could reproduce the mechanical properties of the tissue applying Eq. 4. We took again E’_*N*_ = 0.4kPa and E’_*m*_= 2.0kPa as an initial seed for the fitting procedure [20, 35, 43, 33], assumed that the PNNs have the same stiffness as the NIM, and fitted all values of *E*_*i*_ for each constituent. The fraction, *ρ*_*i*_, was again obtained from the tissue quantification (Fig. 2). We focused on an indentation of *δ* = 13*μ*m (other indentation levels are shown in SI Appendix, Fig. S6-8 and Tables S5-8).

Our theoretical model predicted the experimentally measured mechanical response of all locations in all ROIs with a mean error below 19% (see Fig. 2(l,o) and SI Appendix, Tables S5-6), showing a slight improvement with respect to the affine model (21%). Most regions showed remarkably low errors below 15% except for the GCL (64%), SP3 (26%), SR3 (26%) and SR1 (23%) regions. Note that we obtained these low errors with a clear and physical meaning of the main tissue constituents analyzed in this study. In Table S9, we present the results of E’ for each indentation depth analyzed. The NIM and PNN showed a slightly non-linear strain-stiffening behavior. Considering only one type of ECM resulted in higher errors, which suggests that PNN and NIM could have different stiffness values. The PNN stiffness increased linearly from ∼3kPa, at *δ* = 6.5*μ*m, up to ∼8.5kPa at *δ* = 17*μ*m and the NIM stiffness increased linearly from ∼0.5kPa up to 1.5kPa. Myelin showed a linear increase in the stiffness with increasing *δ* as well (see Fig. S8), from 2kPa up to 7.5kPa. E’ values reported in Table S9 are consistent with directly measured stiffness values of neurons, myelin and ECM [20, 35, 43, 33, 6], which was not achieved in the affine model. Values of the NIM are also consistent with the stiffness of substrates on which neurons and glial cells differentiate [48, 49].

### Viscoelasticity depends on tissue architecture

To investigate whether the additive decomposition of the cSEDF could be used for analyzing not only tissue elasticity but also the viscoelastic behavior of the brain [14, 12], we applied our model to a frequency-dependent characterization of the tissue. We took the loss moduli E”, which represent the viscous component of the mechanical response, of each ROI (see Materials and Methods) and plotted the mean values and standard deviation in Fig. 2(l,n). Then, we applied the relation as in Eq. 4 but for the loss modulus E” to describe the viscoelastic response of each ROI based on its local composition, and performed the same fitting procedure as above (see Table S9). As shown in Fig. 2(l,p), we found a good agreement between experimental data and the theoretical model, with a mean error lower than 16%, and most of the ROIs with errors lower than 15% except, again, the GCL, SP3, and SR3 regions, which slightly overpassed this threshold. The reported loss moduli for all components were again comparable to data from literature [20, 35, 43, 33] and showed similar increases with increasing indentation depths (SI Appendix, Table S9).

### Composition and stiffness of brain components reproduce CNS mechanics

To further investigate the general applicability of our model, we extended our study to the hippocampus of adult mice by using previously reported data [18]. The tissue composition in the adult hippocampus is shown in Fig. 3 and SI Appendix, Table S1. We focused again on *δ* = 13*μ*m (other indentation depths are shown in SI Appendix, Fig. S5 and Tables S3-4,S9). We observed a stiffening of both the ECM (PNN and NIM) and cell bodies compared to the juvenile samples, which is in line with previously published data on rodent brain tissue stiffening during development and ageing [3, 6]. We found a same tendency for the loss modulus E” (see Fig. 3 and SI Appendix, Table S9).

**Figure 3:**
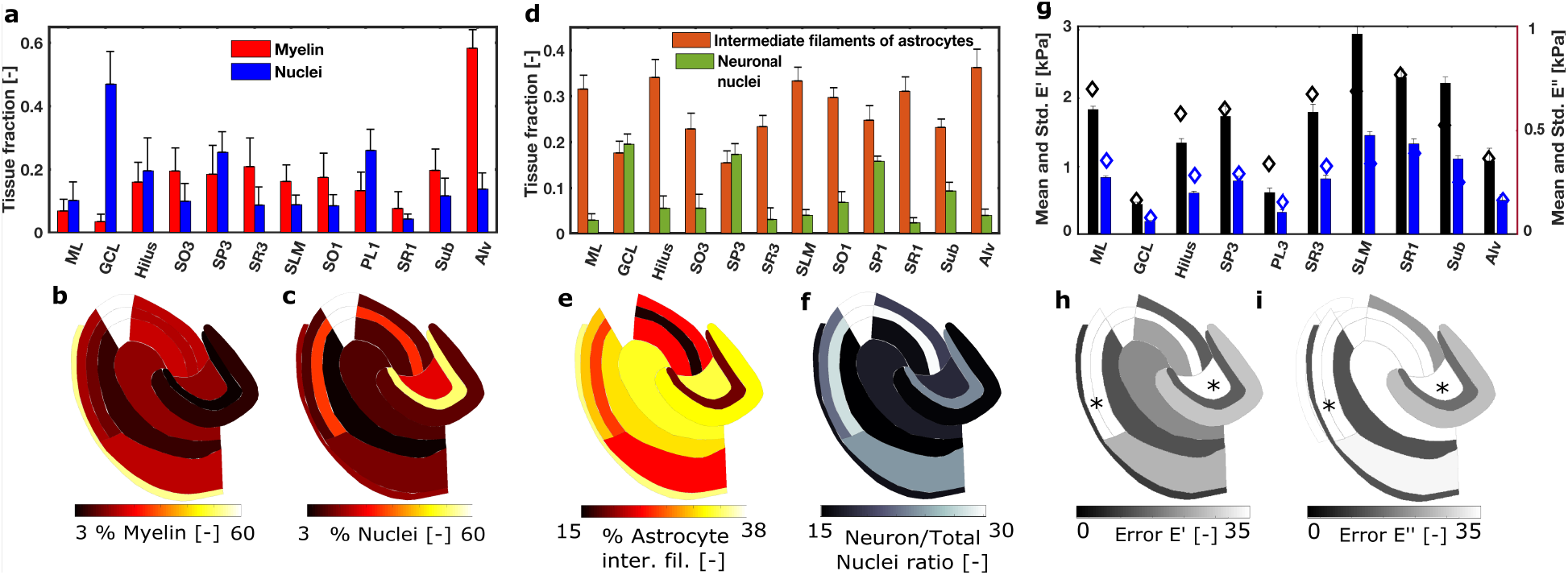
Quantification and analysis of tissue composition and stiffness of ROIs for the hippocampus of juvenile mouse. Mean and standard deviation of the mylein and nuclei (a) and representation of the mean value in each specific ROI (b-c). Mean and standard deviation of the neuron nuclei and intermediate filament of filaments of astrocytes (d) and representation of the mean value of intermedia filaments and the neuron vs. total nuclei in each specific ROI (e-f).Mean and standard deviation of E’ and E” for the different region of the hippocampus and prediction (diamond marks) of the proposed mechanical model (g). Error of the proposed mechanical model for E’ and E” represented in each ROI (h-i).

Using our model (Eq. 4), we were able to reproduce the mechanical properties of the adult hippocampus with an average error of measurements of 25% and 27% for E’ and E”, respectively. Again, these values represent a decrease with respect to the errors obtained with the affine model, 31% and 35% for E’ and E”, respectively. The hilus and SP3 were the only regions with error higher than 40%. It is important to note, however, that the model accurately captured the tendency of the age-related change in tissue rigidity for all ROIs. Moreover, the stiffness values found by the fitting procedure were again within the values reported in literature [20, 35, 43, 33], which was not possible for the affine model. The myelin and NIM stiffness showed a linear increase with increasing indentation depths. However, the myelin stiffness showed a slight decrease with respect to the juvenile samples while the stiffness of the NIM doubled its values with respect to the juvenile animals. The PNN was again the stiffest component of the brain.

Finally, we applied our model to a different region of the brain, the cerebellum of juvenile mice, by using previously reported data [23] (Fig. 4). We predicted E’ and E” of different ROIs (see Fig. 4) considering the distribution of the cerebellum’s components (Fig. 4(d-i)) and the fitted values of E’ obtained for the components of the juvenile hippocampus (Fig. 3 and Table S9). The model was able to reproduce the cerebellum’s heterogenous mechanics [23] (Fig. 4k-j) without any further fitting of parameters and they were again within the values reported in literature [50].

**Figure 4:**
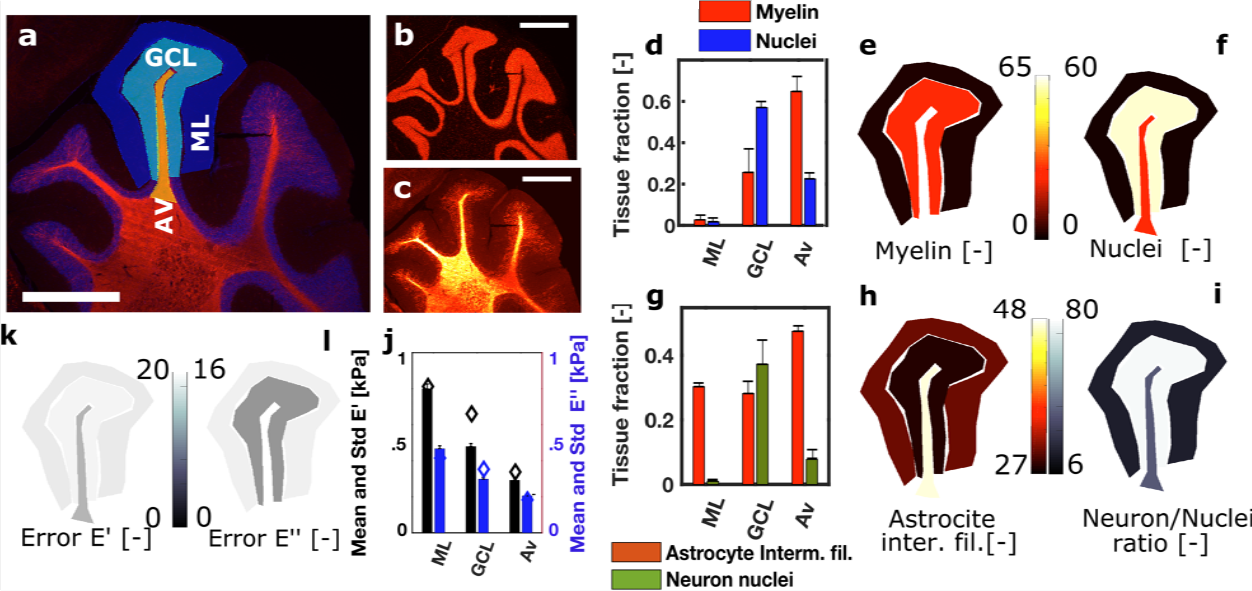
Quantification and analysis of tissue composition and stiffness of ROIs for the cerebellum of juvenile mice (a-c). Nuclei and myelin are shown in blue and red, respectively (a), nuclei (b) and myelin (c). Scale bars (a-e) are 100*μm*. Abbreviations used for regions are: GCL= granular cell layer, ML = molecular layer, Av=Arbor vitae. Mean and standard deviation of tissue brain composition (d-i): Mean and standard deviation of the myelin and nuclei (d) and representation of the mean value in each ROI (e-f). Mean and standard deviation of the neuron nuclei and intermediate filaments of astrocytes (g) and representation of the mean value of filaments and the neuron vs. total nuclei ratio in each specific ROI (h-i). Error of the proposed mechanical model for E’ and E” represented in each ROI (k-l). Mean and standard deviation of E’ and E” for the different region of the cerebellum (j). Diamond marks shows the prediction of the proposed mechanical model.

We found mean errors of 21% and 15% for E’ and E”, respectively, which were lower than the 23% and 41% for E’ and E”, respectively, yielded by the affine model (see also SI Appendix, Fig. S11 and Tables S14-15). In summary, our results indicated that the non-affine model predicts local nervous tissue mechanics well, irrespective of age and regions, solely based on the distribution and mechanical properties of its constituents, and it outperforms the affine model.

## Discussion

In this study, we developed a mathematical model that uses the morphological structure of brain tissue to determine its local mechanical properties of healthy CNS tissue. We found that a classical additive decomposition of the Strain Energy Density Function (SEDF), and equivalently stresses, of each tissue constituent cannot reproduce the complex mechanical behavior of brain tissue. However, when we took the actual structural organization of the tissue into account, we were able to predict brain mechanics. By considering the abundancy and mechanical properties of individual cellular and extracellular tissue components as well as the arrangement of these components within the brain tissue, some being connected in parallel and others in series, we were able to reproduce the local mechanical behavior of the tissue. Finally, we also present an online platform for the scientific community (uploaded in the MATLAB Central File Exchange) that, upon providing immunohistochemistry images and stiffness values for the constituents of the sample, yields the the viscoelastic properties at the tissue scale.

Previously developed models (see [21, 19] for a review) often considered a homogenized response of the tissue constituents to applied forces. Other models have considered different tissue components but not their actual properties, proposing constitutive laws that included parameters with unclear physical interpretation [22, 23] or have focused on one tissue component with stronger correlation with tissue stiffness [24]. Our results suggest that the classical additive decomposition of the SEDF is not appropriate to reproduce tissue stiffness from the structure and stiffness of its components. We argue that this is due to an inherent affine deformation of such models. By incorporating a precise multi-scale description of the tissue, our model now enables us to predict mechanical properties of specific brain tissue regions, which are characterized by distinct compositions and arrangements of cells and ECM.

We followed a bottom-up approach: based on what we knew about the brain tissue constituents, we reproduced local brain tissue mechanics with minimal fitting parameters. Specifically, we only fitted the storage and loss moduli of myelinated axons, perineuronal nets and interstitial matrix. Ideally, this information could be obtained from separate experiments. Here, we fitted their values and made sure they are within the range of data reported in the literature. Furthermore, we used the fitted values of the storage and loss moduli from the juvenile hippocampus (Table S9) and reproduced, without any further fitting procedure, the stiffness of the juvenile cerebellum (Fig. 4). We found that, in juvenile samples, the stiffness of the ECM increases non-linearly with increasing strain, that the stiffness of cell bodies does not change with strain, and that the stiffness of myelin increases with strain. In adult tissue, however, the stiffness to tissue constituents changes variably with changing strain levels. Regions with a high concentration of neurons, such as the GCL or the SP3, were reproduced worse than others. We hypothesize that these regions are more fragile than other regions in the hippocampus and start swelling or even dying earlier, which may explain the observed deviations between data and model.

Our results suggest that there are simple mechanical relationships between tissue constituents that regulate the mechanical properties of brain tissue. Incorporating tissue architecture into models describing tissue mechanics could potentially be used for the constitutive modeling of other soft biological tissues, for example, cardiovascular tissue [25, 40]. There are still important questions to be answered to improve our current understanding of brain tissue mechanics: for example, do neuronal cells and ECM change their mechanical properties during development? How do cell-ECM interactions and adhesive interfaces change during development and ageing? How do the different ECM structures contribute to tissue mechanics? What role does the vasculature and the anisotropy play in regulating brain tissue mechanics?.

Here, we used a viscoelastic model to focus on the arrangement of the tissue constituents. More advanced viscoelastic models could reproduce the behavior of the tissue even more accurately (see, e.g., [51]). Separating the role of viscoelasticity from poroelasticity should also be the focus of research for a complete understanding of brain mechanics. Our model relies on a simple meanfield homogenization method. More advanced homogenization methods (see, e.g., [52, 53]) will require a morphological reconstruction of the representative volumes of interest at high resolution, which is currently the limiting step. Once these data are available, computationally more expensive multiscale finite element methods or advanced mean-field approaches can be applied to investigate brain mechanics with even higher accuracy.

A precise description of how tissue constituents arrange in soft biological tissue will allow us not only to better understand its mechanical properties but also to design better biomedical materials [54] and engineered tissues [55]. The cooperation of mathematical models, computational bioengineering, image analysis and neuroscience will eventually allow us to answer these and many more questions.

## Acknoledgements

P.S has been supported by the Generalitat de Catalunya under grants 2017-SGR-1278. N.A acknowledge funding from the European Research Council (Consolidator award 615170). K.F. acknowledges funding from the European Research Council (Consolidator Award 772426), and the Alexander von Humboldt Foundation for his Alexander von Humboldt Professorship.

## Contributions

P.S designed the work, performed the mechanical simulations. C.B. performed the image analysis. N.A. analyzed the mechanical tests. P.S. and K.F. wrote the paper. All authors analyzed the data and critically reviewed the paper.

## Notes

https://es.mathworks.com/matlabcentral/fileexchange/90531-channelcoloranalysis?s_tid=prof_contriblnk

### Competing Interest Statement

The authors have declared no competing interest.

