## Supplementary material for "Brain tissue mechanics is governed by microscale relations of the tissue constituents": SI Appendix

#### 1 Homogenization method

Different homogenization techniques can be followed based on specific criteria, such as the arrangement of the microstructure, computational cost or required accuracy (see [1, 2, 3] for reviews on homogenization methods). Among them, we can find asymptotic methods, generalized cells, fast Fourier transforms, embedding approaches, mean-field (MF) homogenization schemes and methods based on the Finite Element (FE). The MF and FE-based approaches are the most commonly used strategies to deal with homogenization problems. The FE family of methods is well suited for relatively simple and periodic microstructures because defining a very fine structure may be veryy time consuming and computationally expensive. This is specifically the case in the brain tissue, where the microstructure is extremely complex.

Here, we followed a mean-field (MF) homogenization scheme. MF schemes are easy to implement, cheap to compute and with a good accuracy of the results at the level of the Representative Volume Element (RVE) [4]. Microstructure heterogeneities are treated separately as homogeneous phases, and the quantities of interest in the microstructure are just averaged values for the macroscopic response. The main condition required for the RVE is its statistical representation of the macroscopic material. Therefore, the RVE must be defined such that it contains a large number of microstructural features. Let us consider a macroscopic length scale L and a microscopic length scale d. L must be large enough in comparison to d so that the microscale variables influence the macroscopic behavior through the averaged variables in the microstructure, i.e. d/L¡¡1.

In our problem, we can define the length d at the microstructures that correspond to somas ( 10um), axons ( 1-2um in diameter), and the components of the EMC. We assumed that our RVE, which was on the order of 50um, was large enough compared with the microscopic length scales. Note, that we were limited by the size of some ROIs in the hippocampus, which were as small as 50µm.

#### 1.1 The strain energy function and the affine regime

We started the derivation of our model by introducing the homogenization scheme in a large-strain setting (see, e.g., [5]). First, we defined the Strain Energy Density Functions (SEDF) $\Psi(F)$. The deformation gradient $\mathbf{F} = \nabla_X \varphi(\mathbf{X}, t) \; : T\Omega_0 \to T\Omega_t$ is the linear tangent map from the tangent space $T\Omega_0$ to the time–dependent tangent space $T\Omega_t$. By deriving the SEDF with respect to $\mathbf{F}$, we obtained the thermodynamic force conjugate to $\mathbf{F}$, $\mathbf{P} = \partial_{\mathbf{F}}\Psi$, the first Piola-Kirchhoff stress tensor. All these quantities can be related to either a homogenous macroscopic material or a single microstructural component. In the mean-field theory, a macroscopic value of a quantity $\langle(\bullet)\rangle$ defined in a 3D space $\Omega$ of volume $V$ can be obtained by averaging the microscopic fields $(\bullet)$ over $\Omega$ as

$$\langle(\bullet)\rangle_\Omega = \frac{1}{V}\int_\Omega (\bullet)\,\mathrm{d}V. \tag{1}$$

Based on a standard approach in the modeling of soft biological tissues (e.g. arteries [6, 7], cartilage [8] and brain tissue [9]) and motivated by thermodynamical reasoning, we can obtain the macroscopic or total SEDF $\Psi_T$ of the tissue as

$$\Psi_T = \langle\Psi\rangle_\Omega = \frac{1}{V}\sum_i^n \int_{\Omega_i} \Psi_i\,\mathrm{d}V_i. \tag{2}$$

where we have assumed that the material is made of $n$ components that occupies a 3D space $\Omega_i$ of volume $V_i$. If we define

$$\langle\Psi_i\rangle_{\Omega_i} = \int \Psi_i\,\mathrm{d}V_i \quad \text{then we have that } \Psi_T = \sum_i^n \rho_i \langle\Psi_i\rangle_{\Omega_i}\,, \tag{3}$$

where $\Psi_i$ and $\rho_i$ are the SEDF and volume fraction, respectively, of each tissue constituent $i$.

The derivative of $\Psi_T$ with respect to $\mathbf{F}$ is the macroscopic first Piola-Kirchhoff stress in the tissue:

$$\mathbf{P}_T = \partial_{\mathbf{F}}\Psi_T = \sum_i^n \rho_i \langle\mathbf{P}_i\rangle_{\Omega_i}, \tag{4}$$

where $\langle\mathbf{P}_i\rangle_{\Omega_i}$ are the first Piola-Kirchhoff stress tensor of each tissue constituent. Note that we derived the stress tensor by one single deformation gradient, meaning that all tissue constituents suffer the same deformation. Given the additive decomposition of the SEDF and the single deformation of the tissue, we inherently imposed an affine deformation in the tissue.

Similarly, and within the small strain limit of the experiments, the total Cauchy stress tensor is given as

$$\sigma_T = \sum_i^n \rho_i \sigma_i, \tag{5}$$

where the stress-strain constitutive relation for each constituent is defined as

$$\sigma_i = \mathbb{C}_i : \boldsymbol{\varepsilon} \tag{6}$$

$\boldsymbol{\varepsilon}$ is the infinitesimal strain tensor, or Cauchy's strain tensor, defined as $\boldsymbol{\varepsilon} = \frac{1}{2}\left[\boldsymbol{\nabla}\mathbf{u} + (\boldsymbol{\nabla}\mathbf{u})^{\mathrm{T}}\right]$, where $\mathbf{u}$ is the displacement tensor. Note that, again, we assumed that the strain in each constituent is the same as the macroscopic strain tensor, as discussed above. $\mathbb{C}_i$ is the fourth-order stiffness tensor of each constituent $i$. The constitutive law of the tissue is therefore uniquely determined by the stiffness tensor, which, assuming the tissue to be isotropic and almost-incompressible, is a function of the storage Young's modulus E (a measure of tissue stiffness). Finally, we can convine Eq. 5 and 6 and write $\sigma_T = \sum_i^n \rho_i \mathbb{C}_i : \boldsymbol{\varepsilon}$. Considering the indentation tests, we can write the final stress-strain relation

$$\sigma_T = E_T \varepsilon, \text{ where } E_T = \sum_i^n \rho_i E_i \tag{7}$$

is the total equivalent Young modulus obtained from the indentation tests, that is the total stiffness of the tissue due to the presence of several constituents with Young's moduli $E_i$.

#### 1.2 The complementary strain energy function and the non-affine regime

As we discuss in the main text, the additive decomposition of the SEDF imposes affine deformations in the microstructural constituents because, within this framework, each individual constituent is arranged in parallel and strain is shared equally within all the constituents. However, we know that brain tissue constituents are not only arranged in parallel but also in series (see Fig. 1 and Introduction in the main text). To generalize this idea within a consistent thermodynamic formulation, we took advantage of the complementary energy density function (cSEF) (see [10, 12, 11] among others).

Let us start by recalling the polar decomposition of the deformation gradient as $\mathbf{F} = \mathbf{R} \cdot \mathbf{U}$, where $\mathbf{U}$ is the symmetric positive definite right stretch tensor, defined with respect to the reference configuration. The rotation tensor $\mathbf{R}$ is a unique proper orthogonal tensor such that $\det(\mathbf{R}) = 1$. Following [10], the co-rotated, symmetrized Biot or Jauman material stress tensor $\boldsymbol{\tau}_J$ is defined as

$$2\boldsymbol{\tau}_J = \mathbf{P}\mathbf{R} + \mathbf{R}^T\mathbf{P}^T. \tag{8}$$

such that $\mathbf{P} : \dot{\mathbf{F}} = \boldsymbol{\tau}_J : \dot{\mathbf{U}}$. Because $\boldsymbol{\tau}(\mathbf{U})$ is frame indifferent, we can write that

$$2\boldsymbol{\tau}(\mathbf{U}) = \mathbf{P}(\mathbf{U}) + \mathbf{P}^T(\mathbf{U}). \tag{9}$$

This relation allows us to define $\mathbf{P} = \partial_{\mathbf{F}}\Psi$ and $\boldsymbol{\tau}_J = \partial_{\mathbf{U}}\Psi$.

Finally, and given the equality $\mathbf{P} : \mathbf{F} = \boldsymbol{\tau}_J : \mathbf{U}$, we define a cSEDF $\hat{\Psi}$ such that

$$\Psi + \hat{\Psi} = \mathbf{P} : \mathbf{F} = \boldsymbol{\tau}_J : \mathbf{U}, \tag{10}$$

where $\hat{\Psi}$ is a function of $\mathbf{P}$ or $\boldsymbol{\tau}_J$. It can be shown that $\hat{\Psi}$ possesses frame indifference inherently and, under local invertibility of $\boldsymbol{\tau}_J(\mathbf{U})$, we can express the deformation gradient as $\mathbf{F} = \partial_{\mathbf{P}}\hat{\Psi}$.

Following the same averaging described in the previous section (Eq. 1 and 2), the macroscopic or total cSEDF is given as

$$\hat{\Psi}_T = \sum_i^n \rho_i \langle \hat{\Psi}_i \rangle_{\Omega_i} \text{ where the averaged cSEDF of each constituent, } i, \text{ is} \langle \hat{\Psi}_i \rangle_{\Omega_i} = \int \hat{\Psi}_i \, \mathrm{d}V_i. \tag{11}$$

After deriving with respect to the conjugate stress tensor $\mathbf{P}$, we obtain the total deformation gradient as

$$\mathbf{F}_T = \sum_i^n \partial_{\mathbf{P_i}} \hat{\Psi}_i = \sum_i^n \rho_i \mathbf{F}_i. \tag{12}$$

It is important to note that we make the derivation with respect to a single stress tensor, meaning that we naturally impose that all tissue constituents are going to bear the same stress while their deformation changes. Therefore, the additive decomposition of the cSEDF naturally imposes non-affine deformation gradients in the tissue constituents.

We modified again the previous derivation to consider the linear and small strain regime. We can express the total infinitesimal strain tensor as

$$\boldsymbol{\varepsilon}_T = \sum \rho_i \boldsymbol{\varepsilon}_i, \tag{13}$$

where the stress-strain constitutive relation for each constituent is defined as

$$\boldsymbol{\varepsilon}_i = \hat{\mathbb{C}}_i : \boldsymbol{\sigma}. \tag{14}$$

$\hat{\mathbb{C}}_i$ is the compliance tensor of each tissue constituent, which, again, is a function of E and the Poisson's ratio. Combining the two previous equations, we obtain $\boldsymbol{\varepsilon}_T = \sum_i^n \rho_i \hat{\mathbb{C}}_i : \boldsymbol{\sigma}$. Considering the indentation tests, we can write the final stress-strain relation as

$$\sigma = E_T \varepsilon_T, \text{ where } E_T = \frac{\prod_i^n \rho_i E_i}{\sum_i^n \rho_i E_i} \tag{15}$$

is the total equivalent Young modulus obtained from the indentation tests.

#### 1.3 Relation to classical homogeneization results: the Voigt and Reuss estimates

Classical mean-field theory exactly recapitulates the previous expressions. Within the small strain assumption, the work by Hill [4] showed that the total stress and strain tensors are

$$\boldsymbol{\sigma}_T = \sum \rho_i \boldsymbol{\sigma}_i \text{ and } \boldsymbol{\varepsilon}_T = \sum \rho_i \boldsymbol{\varepsilon}_i, \tag{16}$$

respectively, which are exactly the expressions provided above (see Eq. 7 and 15). These two relations have been shown to fulfills the Hill-Mandel condition, which requires that the averaged mechanical work done within the RVE equals the work computed by using the averaged stress and strain tensor, i.e., $\boldsymbol{\sigma}_T : \boldsymbol{\varepsilon}_T = \langle \boldsymbol{\sigma} : \boldsymbol{\varepsilon} \rangle_{\Omega_i}$. Therefore, the expressions derived in the previous sections fulfills the Hill-Mandel condition and allow us to consistently homogenize across scales.

Next, constitutive relations for each tissue constituent have to be provided. In the linear and small strain regime, we can express this relation as $\sigma_i = \mathbb{C}_i : \boldsymbol{\varepsilon}$. However, the stress and strain relations for every constituent must be given so that the stress or the strain, respectively, can be computed, otherwise the problem is not closed. The general form of these expressions are generally given as

$$\boldsymbol{\sigma}_i = \mathbb{A}_i : \boldsymbol{\sigma}_T \text{ and } \boldsymbol{\varepsilon}_i = \mathbb{B}_i : \boldsymbol{\varepsilon}_T, \tag{17}$$

where $\mathbb{A}$ and $\mathbb{B}$ are known as the fourth-order stress and strain concentration tensor, respectively. Different homogenization schemes arise from the choice of these tensors, such as the Mori-tanaka model, the self-consistent scheme or the Eshelby's solution, among many others. Among them, the Voigt and Reuss schemes are the two most simple approaches [13, 14, 15] and, they represent upper and lower bounds of the averaged macroscopic response. The Voigt scheme assumes that $\mathbb{A} = \mathbb{I}$ and, therefore, it assumes affines deformations while the stress varies within the microconstituents. $\mathbb{I}$ represents the fourth order identity tensor. On the other hand, the Reuss scheme assumes that $\mathbb{B} = \mathbb{I}$ and, therefore, imposes the same stress and allows the strain to vary for each constituent. In conclusion, the affine and non-affine models derived from the SEDF and cSEDF above are equivalent to the Voigt and Reuss estimates from classical mean-field homogenization schemes.

Some models in the MF approach, such as the Mori-Tanaka model or the self-consistent approach, require to solve the interaction of the subdomains, usually by analytical solutions of a simplified microstructural problem, such that the averaged quantities fulfill the boundary conditions.

Here, we followed a simple mean-field analysis based on Voigt and Reuss estimates [13, 14, 15], which do not require a solution of the boundary value problem at the microscale and assume perfect bonding between constituents, i.e. no slip conditions.

#### 2 Sample preparation and ROI identification

All experimental protocols have been previously described and published (see details in [16, 17] and Supporting Information). In summary, horizontal brain tissue slices of 300$\mu$m thickness were extracted from 3 to 4 mm of dorsal-ventral positions from two age groups of wild-type mice (C57Bl6/Harlen): juvenile (1 month- old) and young adult (6 and 9-month-old). Each brain tissue slice was placed in a glass bottom chamber for imaging with an inverted microscope, supplied with carbogenated artificial cerebrospinal fluid to maintain the viability and gently pressed down with harp for stabilization. All measurements were performed within 8 hours. Up to 12 regions of interest were identified in the hippocampus and three in the cerebellum [16, 17].

#### 3 Indentation protocol

We refer to [17] for further details on the testing protocol. In summary, the indentation setup consisted of a cantilever-based ferrule-top probe (0.2-0.5 N/m stiffness and 60-105$\mu$m bead radius) mounted on a piezo transducer and XYZ micromanipulator. Indentation was performed by imposing oscillations of 0.2$\mu$m amplitude and 5.62 Hz frequency on top of the ramp loading at 0.01 strain rate in an indentation-depth controlled mode. The raw data was analyzed with an in-house Matlab code to obtain the storage and loss moduli:

$$E'(\omega) = \frac{F_0}{h_0} cos(\delta) \frac{\sqrt{\pi}(1-\nu^2)}{2\sqrt{A}} \tag{18}$$

$$E"(\omega) = \frac{F_0}{h_0} sin(\delta) \frac{\sqrt{\pi}(1-\nu^2)}{2\sqrt{A}} \tag{19}$$

where $F_0$ and $h_0$ are the amplitudes of oscillatory load and indentation-depth, respectively, $\delta$ is the phase-shift between indentation and load oscillations, $\nu$ is the Poisson's ratio (assumed to be 0.5 as brain tissue is a nearly incompressible material) and $A$ is the contact area. Indentation mapping was performed at 50$\mu$m step size over regions in hippocampus and cerebellum for tissue slices from juvenile brain (N=6) and only over hippocampus for adult brain slices (N=5). Each indentation location was assigned to the measured ROI.

Nanoindentation is a classical approach to measure the mechanics of cells and tissues at the micro and nano scale. Nanoindentation has been widely used to test soft brain tissue [18, 19, 20, 21, 17, 22, 23, 16] as well as in cell mechanics (see, e.g., [24, 25]. The indentation is usually controlled such that the small strain limit is ensured based on classical considerations. Specifically, indentation displacements were imposed on the tissue such that $\epsilon < 0.05$. Following the Hertz model, the contact radius $a = \sqrt{h}R$ varied between 22–39 $\mu$m in order to fulfill the small strain approximation $\epsilon < 0.08$ [26], where $h$ is the indentation depth and $R$ is radius of the indentater tip.

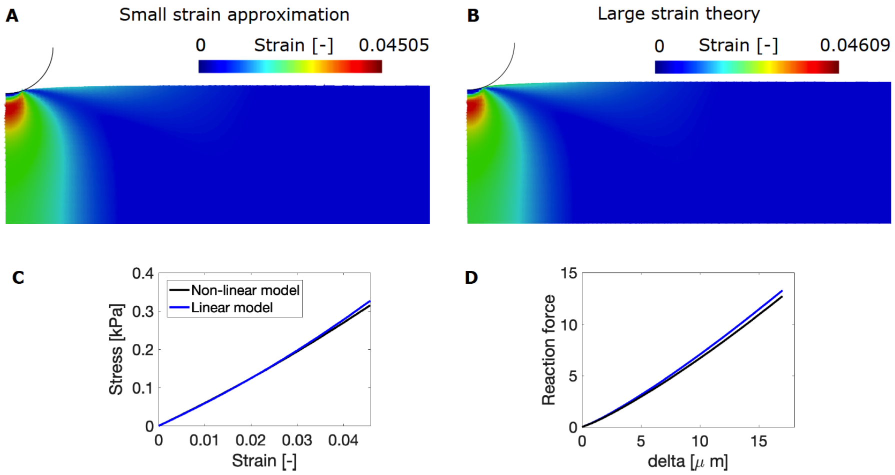


Figure 1: Finite element simulation of the indentation test performed in our experiments. The thickness of the sample is 300$\mu$m, the bead radius is 105$\mu$m and we indent 17$\mu$m so that $a = 42.24\mu$m and $\epsilon = 0.08$. (A-B) We show the maximum principal strain for the small strain theory and the large strain models with a Neo-Hookean constitutive law, respectively. (C) Stress-train relation for the small and large strain theory. (D) Reaction force-indetation depth for the small and large strain theory.

To analyze how good the small strain approximation is compared to the large strain theory, we performed 2 FE models (see Fig. 1) in the FE software Abaqus. The first model was run based on the small strain theory (Fig. 1.A) while the second (Fig. 1.B) was computed following large strain and a Neo-Hookean ($\Psi = C(I_1 - 3)$) strain theory. To compare both models, we assumed a Young's modulus of E=3kPa and an equivalent material parameter C=0.5kPa. Both material parameters are related such that 6C=E. As we can see in Fig. 1, where we plot the maximum principal strain, both approaches behave very similarly in the strain regime of the experiments, with maximum strains of 4.609% for the large strain theory (Fig. 1.B) and 4.505% for the small strain model (Fig. 1.A). We also plot the stress-strain relation (Fig. 1.C) and the reaction force as a function of the indentation depth (Fig. 1.D). Similarly, both approaches showed a very similar behavior, suggesting that the small strain regime is valid in this experimental approach. which has been taken by many others in the past [18, 19, 20, 21, 17, 22, 23, 16, 24, 25].

### 4 Image adquisition

Images of immunohistochemically stained slices were previously obtained and used here for further analysis [16]. In short, two age groups of 3 wild-type mice (C57Bl6/J) were used to stain nuclei of neurons (NeuN), all cell nuclei (Hoechst), astrocytes (GFAP), myelin (MBP) and dendrites (MAP2). Details on the antibodies used for the (immuno)histochemical staining are provided elsewhere [16]. Fluorescence images were obtained with a Zeiss Axioscope.A1 epi-microscope with a 10x Plan-NeoFluar objective. Anatomical regions were identified and drawn manually as polygons using ImageJ and their coordinates stored in .roi files.

### 5 Density quantification for all the samples

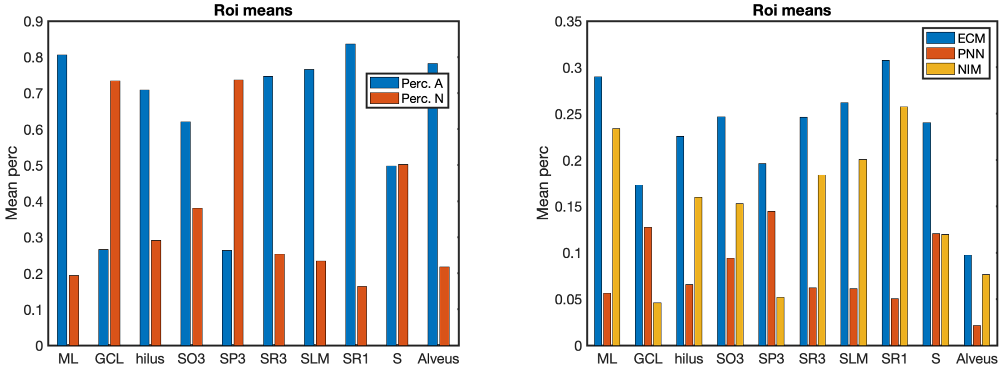


Figure 2: Percentage of astrocytes and neurons of all cell nuclei (left) and percentage of ECM, PNN and NIM (right) obtained as described in the main text for the samples of the adult hippocampus.

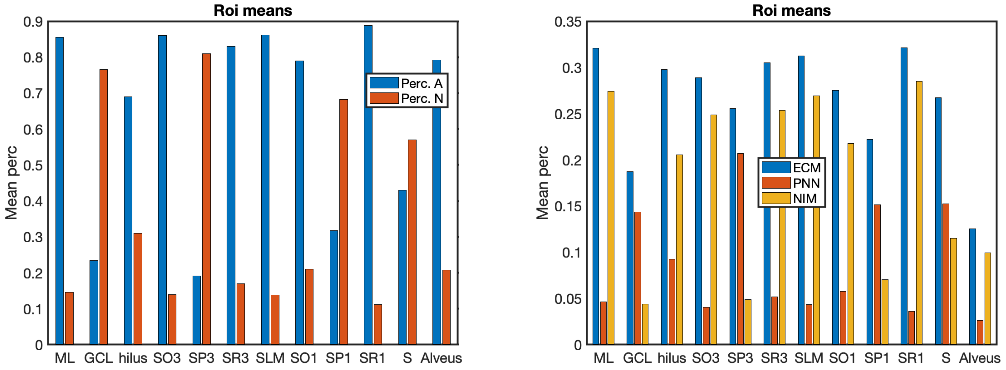


Figure 3: Percentage of astrocytes and neurons of all cell nuclei (left) and percentage of ECM, PNN and NIM (right) obtained as described in the main text for the samples of the juvenile hippocampus.

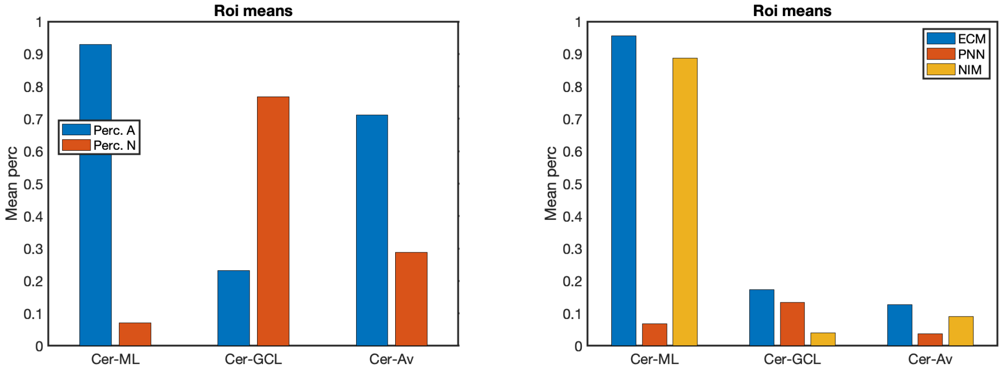


Figure 4: Percentage of astrocytes and neurons of all cell nuclei (left) and percentage of ECM, PNN and NIM obtained as described in the main text (right) for the samples of the juvenile cerebellum.

| Region | - | ML | GCL | hilus | SO3 | SP3 | SR3 | SLM | SO1 | SP1 | SR1 | S | Alveus/Av |
|---|---|---|---|---|---|---|---|---|---|---|---|---|---|
| Juvenile Hippo. | $\rho$ Myelin [-] | 0.017 | $5.3\,10^{-3}$ | 0.032 | 0.12 | 0.083 | 0.09 | 0.035 | 0.13 | 0.11 | 0.036 | 0.11 | 0.5 |
| | std [-] | $4.0\,10^{-3}$ | $4.5\,10^{-3}$ | 0.016 | 0.069 | 0.07 | 0.058 | 0.019 | 0.077 | 0.072 | 0.03 | 0.06 | 0.11 |
| | $\rho$ Nuclei [-] | 0.067 | 0.46 | 0.12 | 0.052 | 0.19 | 0.038 | 0.072 | 0.082 | 0.25 | 0.046 | 0.12 | 0.15 |
| | std [-] | 0.012 | $9.6\,10^{-3}$ | 0.031 | 0.011 | 0.043 | $9.8\,10^{-3}$ | $7.2\,10^{-3}$ | 0.016 | 0.02 | $8.3\,10^{-3}$ | 0.026 | 0.025 |
| | $\rho$ Astro. [-] | 0.27 | 0.17 | 0.33 | 0.23 | 0.11 | 0.22 | 0.35 | 0.26 | 0.23 | 0.3 | 0.22 | 0.43 |
| | std [-] | 0.026 | 0.039 | 0.042 | 0.031 | 0.035 | 0.031 | 0.046 | 0.031 | 0.023 | 0.027 | 0.026 | 0.055 |
| | $\rho$ Neurons [-] | 0.019 | 0.33 | 0.059 | 0.015 | 0.28 | 0.018 | 0.022 | 0.028 | 0.2 | 0.015 | 0.12 | 0.045 |
| | std [-] | 0.011 | 0.02 | 0.036 | 0.012 | 0.044 | 0.012 | 0.018 | 0.015 | 0.025 | $7.2\,10^{-3}$ | 0.028 | 0.035 |
| Adult Hippo. | $\rho$ Myelin [-] | 0.069 | 0.036 | 0.16 | 0.2 | 0.19 | 0.21 | 0.16 | - | - | 0.077 | 0.2 | 0.58 |
| | std [-] | 0.037 | 0.023 | 0.062 | 0.073 | 0.091 | 0.09 | 0.052 | - | - | 0.053 | 0.067 | 0.059 |
| | $\rho$ Nuclei | 0.1 | 0.47 | 0.2 | 0.1 | 0.25 | 0.088 | 0.089 | - | - | 0.044 | 0.12 | 0.14 |
| | std [-] | 0.058 | 0.1 | 0.1 | 0.056 | 0.065 | 0.057 | 0.03 | - | - | 0.016 | 0.056 | 0.051 |
| | $\rho$ Astro. | 0.32 | 0.14 | 0.34 | 0.23 | 0.12 | 0.23 | 0.33 | - | - | 0.31 | 0.23 | 0.36 |
| | std [-] | 0.03 | 0.025 | 0.039 | 0.034 | 0.026 | 0.024 | 0.03 | - | - | 0.031 | 0.018 | 0.04 |
| | $\rho$ Neurons | 0.03 | 0.23 | 0.056 | 0.056 | 0.21 | 0.032 | 0.041 | - | - | 0.024 | 0.094 | 0.04 |
| | std [-] | 0.014 | 0.022 | 0.027 | 0.03 | 0.023 | 0.025 | 0.012 | - | - | 0.011 | 0.019 | 0.014 |
| Juvenile Cer. | $\rho$ Myelin [-] | 0.027 | 0.256 | - | - | - | - | - | - | - | - | - | 0.648 |
| | std [-] | 0.022 | 0.114 | - | - | - | - | - | - | - | - | - | 0.072 |
| | $\rho$ Nuclei | 0.017 | 0.57 | - | - | - | - | - | - | - | - | - | 0.225 |
| | std [-] | 0.019 | 0.029 | - | - | - | - | - | - | - | - | - | 0.029 |
| | $\rho$ Astro. | 0.302 | 0.281 | - | - | - | - | - | - | - | - | - | 0.474 |
| | std [-] | 0.011 | 0.037 | - | - | - | - | - | - | - | - | - | 0.016 |
| | $\rho$ Neurons | 0.009 | 0.372 | - | - | - | - | - | - | - | - | - | 0.079 |
| | std [-] | 0.034 | 0.074 | - | - | - | - | - | - | - | - | - | 0.027 |

Table 1: Quantification (mean and std) of myelin, all cell nuclei, intermediate filaments of astrocytes and nuclei of neurons for each ROI in the adult hippocampus, juvenile hippocampus and juvenile cerebellum.

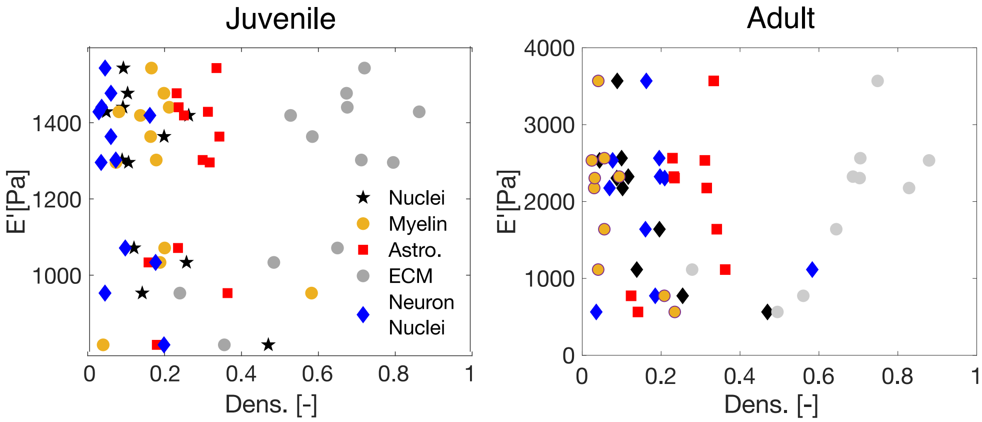


Figure 5: Correlation of stiffness and density of constituents for the juvenile (left) and adult (right) samples of the hippocampus. Negative correlation of all tissue constituents except for the density of intermediate filaments of astrocytes in juvenile and adult mice are observed. The correlation between the stiffness of the tissue and the density of total nuclei and neuronal nuclei shown a significant Pearson coefficient (r=-0.69 and r=-0.67, respectively) for the juvenile mice. For the adult mice, a significant Pearson coefficient for the total nuclei, neuronal nuclei and intermediate filament of astrocytes was found (-0.83, -0.73 and 0.63, respectively). Same behavior is observed for E".

| | Juvenile | | | | Adult | | | |
|---|---|---|---|---|---|---|---|---|
| | $\rho$ Myelin [-] | $\rho$ Nuclei. [-] | $\rho$ Astro. [-] | $\rho$ Neurons [-] | $\rho$ Myelin [-] | $\rho$ Nuclei. [-] | $\rho$ Astro. [-] | $\rho$ Neurons [-] |
| E' | -0.33 | -0.69 | 0.23 | -0.66 | 0.24 | -0.83 | 0.63 | -0.73 |
| E" | -0.37 | -0.58 | 0.20 | -0.57 | 0.29 | -0.83 | 0.672 | -0.76 |

Table 2: Quantification (mean and std) of myelin, all cell nuclei, intermediate filaments of astrocytes and nuclei of neurons for each ROI in the adult hippocampus, juvenile hippocampus and juvenile cerebellum.

### 6 Analysis of E' and E" in adult samples of the hippocampus: Non-affine model

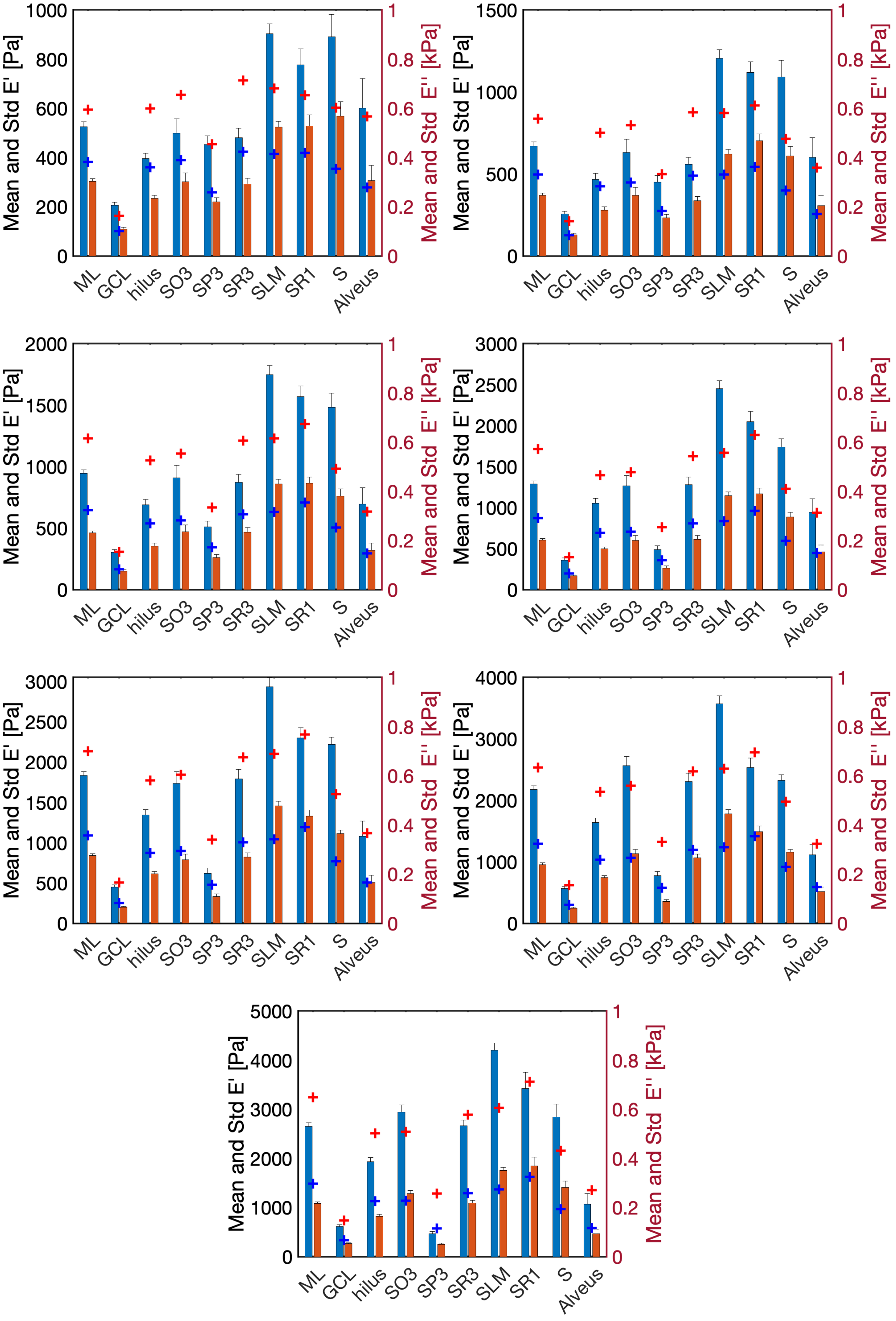


Figure 6: Stiffness values at different ROIs of adult mice. Figures show from top to bottom and left to right the results for 7 indentation strains: 5, 5.5, 6, 6.5, 7, 7.5 and 8%, respectively. Mean and standard deviation of E' and E" for the different regions of the hippocampus. + marks shows the prediction of the proposed mechanical model.

| $0.2\sqrt{R\delta}$/R | ML | GCL | hilus | SO3 | SP3 | SR3 | SLM | SR1 | S | Alveus | Mean per strain |
|---|---|---|---|---|---|---|---|---|---|---|---|
| 5.0 | 0.13 | 0.21 | 0.51 | 0.31 | 1.6e-3 | 0.49 | 0.25 | 0.16 | 0.32 | 0.057 | 0.24 |
| 5.5 | 0.25 | 0.17 | 0.61 | 0.27 | 0.11 | 0.57 | 0.28 | 0.18 | 0.35 | 0.1 | 0.29 |
| 6.0 | 0.3 | 0.016 | 0.52 | 0.21 | 0.31 | 0.39 | 0.3 | 0.14 | 0.34 | 0.088 | 0.26 |
| 6.5 | 0.33 | 0.1 | 0.32 | 0.13 | 0.56 | 0.27 | 0.32 | 0.079 | 0.29 | 1.9e-3 | 0.24 |
| 7.0 | 0.16 | 0.13 | 0.32 | 0.061 | 0.68 | 0.15 | 0.28 | 0.017 | 0.28 | 0.034 | 0.21 |
| 7.5 | 0.17 | 0.11 | 0.3 | 0.13 | 0.71 | 0.073 | 0.29 | 0.096 | 0.15 | 0.16 | 0.22 |
| 8.0 | 0.23 | 0.19 | 0.3 | 0.14 | 1.7 | 0.084 | 0.28 | 0.04 | 0.24 | 0.27 | 0.35 |
| Mean per ROI | 0.22 | 0.13 | 0.41 | 0.18 | 0.59 | 0.29 | 0.29 | 0.1 | 0.28 | 0.1 | |

Table 3: Error of the model's prediction for E' in the adult hippocampus.

| $0.2\sqrt{R\delta}$/R | ML | GCL | hilus | SO3 | SP3 | SR3 | SLM | SR1 | S | Alveus | Mean per strain |
|---|---|---|---|---|---|---|---|---|---|---|---|
| 5.0 | 0.26 | 0.069 | 0.54 | 0.29 | 0.18 | 0.45 | 0.21 | 0.21 | 0.38 | 0.09 | 0.27 |
| 5.5 | 0.35 | 0.027 | 0.53 | 0.21 | 0.18 | 0.45 | 0.2 | 0.23 | 0.35 | 0.17 | 0.27 |
| 6.0 | 0.4 | 0.099 | 0.52 | 0.2 | 0.32 | 0.31 | 0.26 | 0.18 | 0.34 | 0.074 | 0.27 |
| 6.5 | 0.45 | 0.17 | 0.4 | 0.17 | 0.36 | 0.31 | 0.27 | 0.18 | 0.33 | 0.031 | 0.27 |
| 7.0 | 0.29 | 0.28 | 0.42 | 0.13 | 0.45 | 0.23 | 0.28 | 0.1 | 0.31 | 1.0e-3 | 0.25 |
| 7.5 | 0.35 | 0.23 | 0.39 | 0.061 | 0.64 | 0.12 | 0.31 | 0.05 | 0.21 | 0.15 | 0.25 |
| 8.0 | 0.37 | 0.28 | 0.37 | 0.11 | 1.3 | 0.18 | 0.22 | 0.12 | 0.31 | 0.25 | 0.35 |
| Mean per ROI | 0.35 | 0.17 | 0.45 | 0.17 | 0.49 | 0.29 | 0.25 | 0.15 | 0.32 | 0.11 | |

Table 4: Error of the model's prediction for E” in the adult hippocampus.

### 7 Analysis of E' and E" in juvenile samples of the hippocampus: Non-affine model

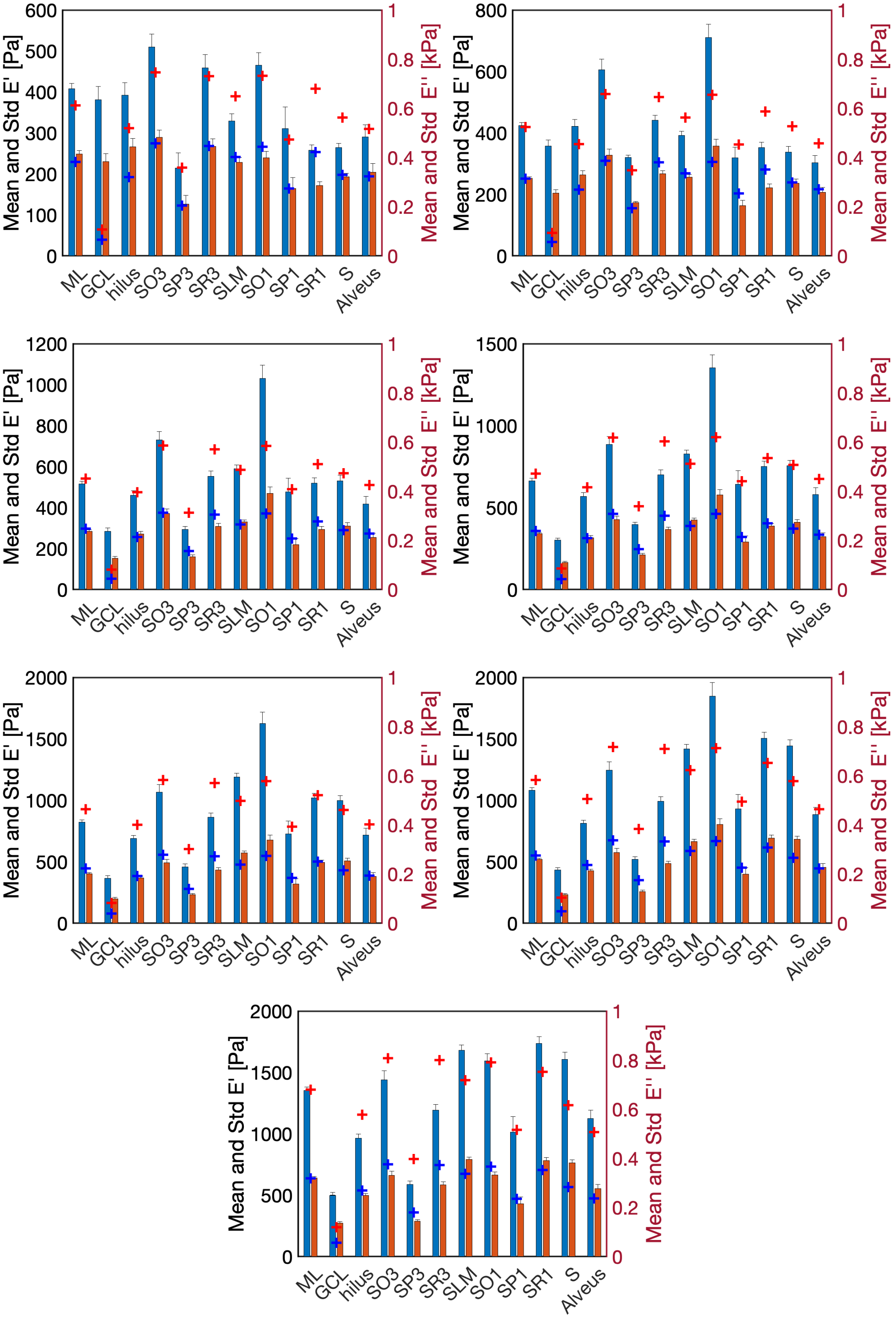


Figure 7: Stiffness values at different ROIs of juvenile mice. Figures show from top to bottom and left to right the results for 7 indentation strains: 5, 5.5, 6, 6.5, 7, 7.5 and 8%, respectively. Mean and standard deviation of E' and E" for the different regions of the hippocampus. + marks shows the prediction of the proposed mechanical model.

| $0.2\sqrt{R\delta}$/R | ML | GCL | hilus | SO3 | SP3 | SR3 | SLM | SO1 | SP1 | SR1 | S | Alveus | Mean per strain |
|---|---|---|---|---|---|---|---|---|---|---|---|---|---|
| 5.0 | 0.1 | 0.83 | 0.2 | 0.12 | 7.9e-3 | 0.044 | 0.18 | 0.056 | 0.086 | 0.58 | 0.28 | 0.069 | 0.21 |
| 5.5 | 0.012 | 0.79 | 0.14 | 0.13 | 0.13 | 0.17 | 0.15 | 0.26 | 0.14 | 0.34 | 0.25 | 0.21 | 0.23 |
| 6.0 | 0.051 | 0.66 | 0.029 | 0.038 | 0.28 | 0.24 | 8.5e-3 | 0.32 | 0.027 | 0.18 | 0.069 | 0.22 | 0.18 |
| 6.5 | 0.066 | 0.58 | 0.098 | 0.046 | 0.28 | 0.29 | 0.074 | 0.31 | 0.027 | 0.066 | 6.9e-3 | 0.16 | 0.17 |
| 7.0 | 0.13 | 0.55 | 0.16 | 0.092 | 0.31 | 0.32 | 0.16 | 0.29 | 0.08 | 0.021 | 0.08 | 0.12 | 0.19 |
| 7.5 | 0.077 | 0.52 | 0.25 | 0.15 | 0.48 | 0.43 | 0.12 | 0.23 | 0.061 | 0.13 | 0.2 | 0.05 | 0.22 |
| 8.0 | 4.6e-3 | 0.52 | 0.2 | 0.12 | 0.35 | 0.34 | 0.14 | 6.8e-3 | 0.019 | 0.13 | 0.23 | 0.097 | 0.18 |
| Mean per ROI | 0.063 | 0.64 | 0.15 | 0.1 | 0.26 | 0.26 | 0.12 | 0.21 | 0.062 | 0.21 | 0.16 | 0.13 | |

Table 5: Error of the model's prediction for E' in the juvenile hippocampus.

| $0.2\sqrt{R\delta}$/R | ML | GCL | hilus | SO3 | SP3 | SR3 | SLM | SO1 | SP1 | SR1 | S | Alveus | Mean per strain |
|---|---|---|---|---|---|---|---|---|---|---|---|---|---|
| 5.0 | 0.081 | 0.83 | 0.28 | 0.048 | 0.027 | 6.2e-3 | 0.058 | 0.12 | 3.4e-3 | 0.47 | 0.022 | 0.047 | 0.17 |
| 5.5 | 2.3e-3 | 0.78 | 0.18 | 0.053 | 0.11 | 0.14 | 0.047 | 0.14 | 0.25 | 0.26 | 9.4e-3 | 0.044 | 0.17 |
| 6.0 | 0.04 | 0.66 | 0.061 | 9.1e-3 | 0.17 | 0.19 | 0.038 | 0.21 | 0.13 | 0.13 | 0.065 | 0.072 | 0.15 |
| 6.5 | 0.049 | 0.61 | 0.012 | 0.085 | 0.16 | 0.23 | 0.089 | 0.2 | 0.1 | 0.044 | 0.092 | 0.035 | 0.14 |
| 7.0 | 0.11 | 0.6 | 0.041 | 0.13 | 0.21 | 0.26 | 0.17 | 0.19 | 0.15 | 0.012 | 0.15 | 9.7e-3 | 0.17 |
| 7.5 | 0.062 | 0.58 | 0.11 | 0.17 | 0.35 | 0.37 | 0.12 | 0.17 | 0.13 | 0.11 | 0.22 | 0.024 | 0.2 |
| 8.0 | 3.6e-3 | 0.59 | 0.081 | 0.14 | 0.25 | 0.27 | 0.15 | 0.1 | 0.088 | 0.098 | 0.26 | 0.14 | 0.18 |
| Mean per ROI | 0.049 | 0.66 | 0.11 | 0.091 | 0.18 | 0.21 | 0.095 | 0.16 | 0.12 | 0.16 | 0.12 | 0.053 | |

Table 6: Error of the model's prediction for E" in the juvenile hippocampus.

### 8 Analysis of E’ and E” in juvenile samples of the cerebellum: Non-affine model

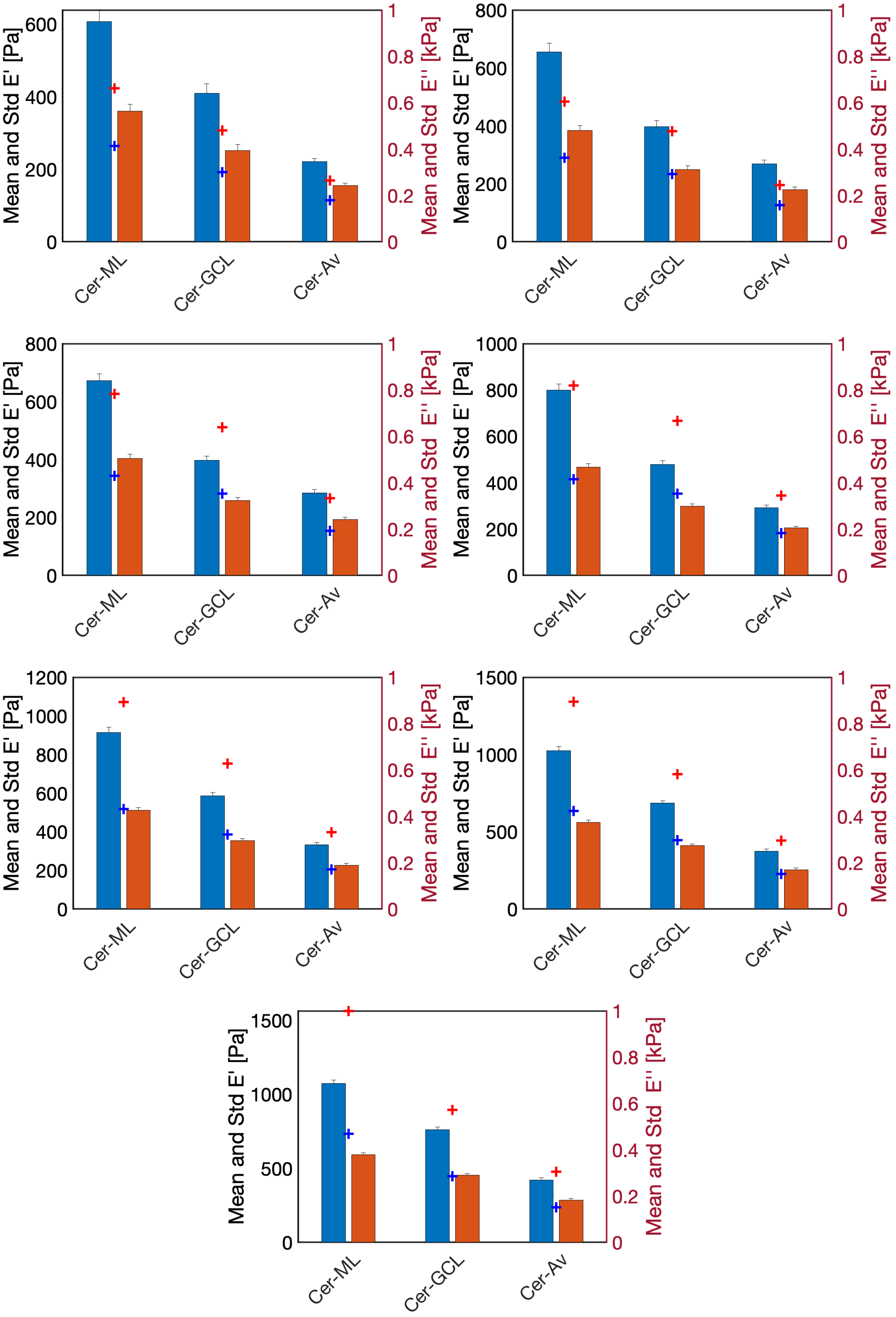


Figure 8: Stiffness values at different ROIs of the cerebellum of juvenile mice. Figures show from top to bottom and left to right the results for 7 indentation strains: 5, 5.5, 6, 6.5, 7, 7.5 and 8%, respectively. Mean and standard deviation of E’ and E” for the different regions of the hippocampus. + marks shows the prediction of the proposed mechanical model.

| $0.2\sqrt{R\delta}$/R | ML | GCL | Av | Mean per strain |
|---|---|---|---|---|
| 5.0 | 0.3 | 0.25 | 0.24 | 0.26 |
| 5.5 | 0.26 | 0.041 | 0.27 | 0.19 |
| 6.0 | 0.067 | 0.29 | 0.064 | 0.14 |
| 6.5 | 0.025 | 0.39 | 0.18 | 0.2 |
| 7.0 | 0.17 | 0.28 | 0.2 | 0.22 |
| 7.5 | 0.31 | 0.27 | 0.19 | 0.26 |
| 8.0 | 0.46 | 0.17 | 0.13 | 0.26 |
| Mean per ROI | 0.23 | 0.24 | 0.18 | |

Table 7: Error of the model's prediction for E' in the juvenile cerebellum.

| $0.2\sqrt{R\delta}$/R | ML | GCL | Av | Mean per strain |
|---|---|---|---|---|
| 5.0 | 0.27 | 0.24 | 0.26 | 0.25 |
| 5.5 | 0.24 | 0.063 | 0.3 | 0.2 |
| 6.0 | 0.15 | 0.091 | 0.2 | 0.15 |
| 6.5 | 0.11 | 0.18 | 0.11 | 0.13 |
| 7.0 | 0.011 | 0.086 | 0.098 | 0.065 |
| 7.5 | 0.13 | 0.084 | 0.11 | 0.11 |
| 8.0 | 0.24 | 0.016 | 0.17 | 0.14 |
| Mean per ROI | 0.17 | 0.11 | 0.18 | |

x

Table 8: Error of the model's prediction for E' in the juvenile cerebellum.

### 9 Changes in E' and E" along indentation strain values in all samples: Non-affine model

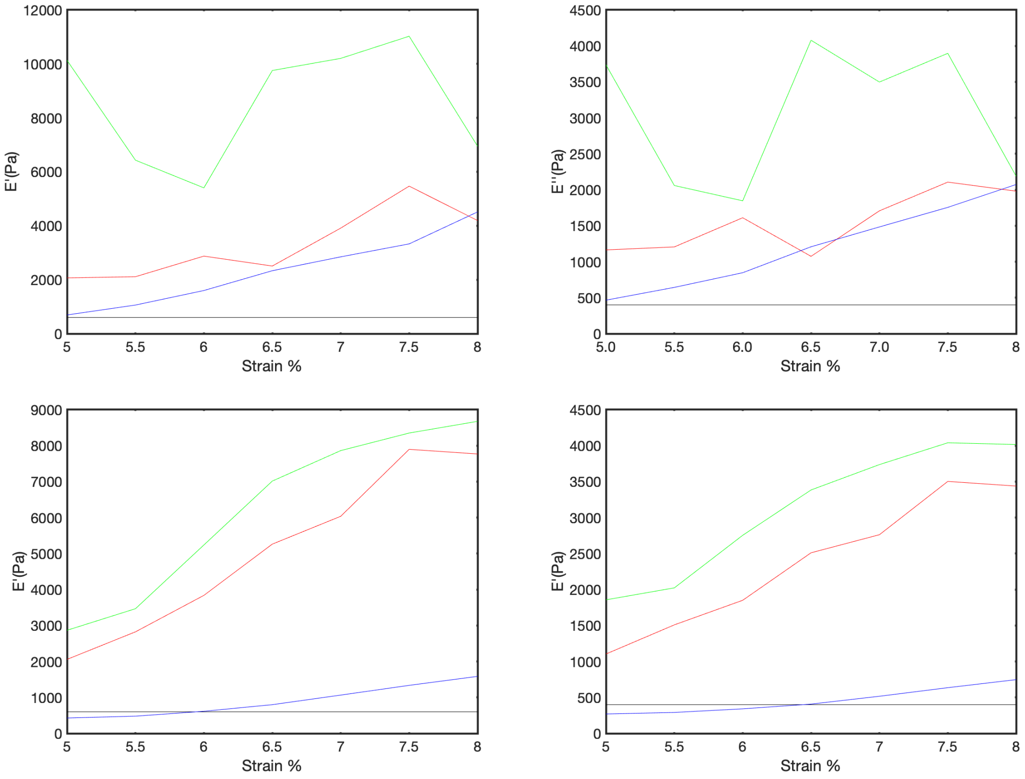


Figure 9: Evolution of E' (left) and E" (right) for adult (top) and juvenile mice (bottom) as a function of the indentation strains, 5.5, 6, 6.5, 7, 7.5 and 8%, of the proposed model. Black, red, green and blue curves for cell bodies, myelin, PNN and NIM, respectively.

Table 9: Predicted values of E' and E" for the components of the mice hippocampus for the juvenile and adult samples.

| | Regios | 5% | 5.5% | 6% | 6.5% | 7% | 7.5% | 8% |
|---|---|---|---|---|---|---|---|---|
| Juv. | | | | | | | | |
| E' [kPa] | Nuclei | 0.6 | 0.6 | 0.6 | 0.6 | 0.6 | 0.6 | 0.6 |
| | Myelin | 2.05 | 2.82 | 3.83 | 5.26 | 6.03 | 7.89 | 7.76 |
| | PNN | 2.86 | 3.46 | 5.23 | 7.01 | 7.86 | 8.35 | 8.67 |
| | NIM | 0.27 | 0.47 | 0.61 | 0.8 | 1.06 | 1.33 | 1.58 |
| Juv. | | | | | | | | |
| E" [kPa]. | Nuclei | 0.4 | 0.4 | 0.4 | 0.4 | 0.4 | 0.4 | 0.4 |
| | Myelin | 1.1 | 1.5 | 1.85 | 2.51 | 2.76 | 3.5 | 3.43 |
| | PNN | 1.85 | 2.02 | 2.75 | 3.38 | 3.73 | 4.04 | 4.01 |
| | NIM | 0.27 | 0.29 | 0.34 | 0.4 | 0.51 | 0.63 | 0.74 |
| Adult | | | | | | | | |
| E' [kPa] | Nuclei | 0.6 | 0.6 | 0.6 | 0.6 | 0.6 | 0.6 | 0.6 |
| | Myelin | 2.07 | 2.11 | 2.87 | 2.5 | 3.92 | 5.53 | 4.19 |
| | PNN | 10.01 | 6.43 | 5.4 | 9.77 | 10.02 | 11.02 | 6.94 |
| | NIM | 0.69 | 1.06 | 1.59 | 2.33 | 2.84 | 3.32 | 4.51 |
| Adult | | | | | | | | |
| E" [kPa]. | Nuclei | 0.4 | 0.4 | 0.4 | 0.4 | 0.4 | 0.4 | 0.4 |
| | Myelin | 1.16 | 1.2 | 1.61 | 1.07 | 1.7 | 2.1 | 1.98 |
| | PNN | 0.84 | 1.15 | 1.41 | 2.11 | 2.59 | 2.81 | 3.86 |
| | NIM | 0.46 | 0.64 | 0.84 | 1.2 | 1.48 | 1.75 | 2.07 |

#### 10 Predicted E’ and E” along indentation strain values in juvenile and adult samples: Non-affine model

#### 11 Analysis of E’ and E” in adult samples of the hippocampus: Affine model

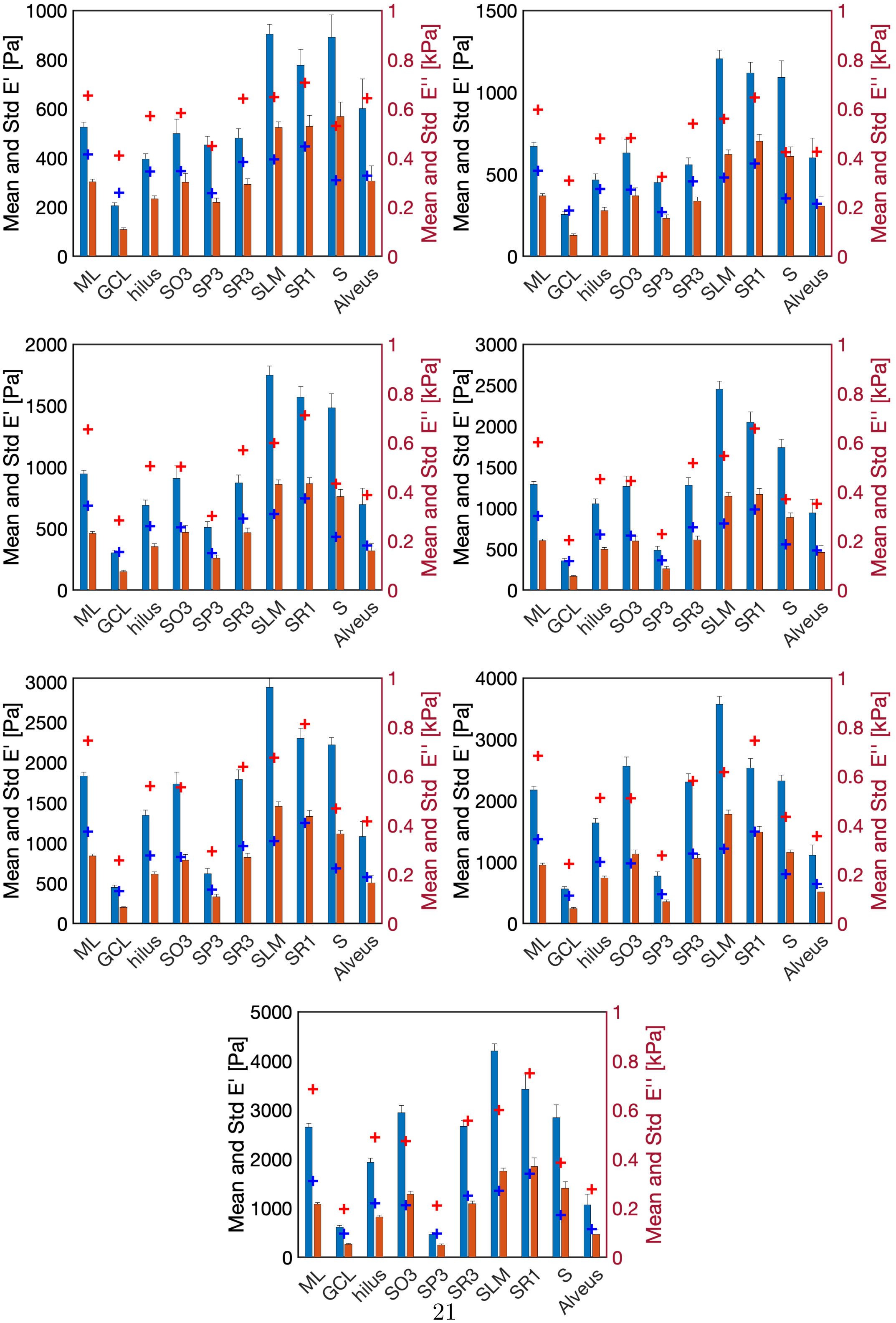




Figure 10: Stiffness values at different ROIs of adult mice. Figures show from top to bottom and

| $0.2\sqrt{R\delta}$/R | ML | GCL | hilus | SO3 | SP3 | SR3 | SLM | SR1 | S | Alveus | Mean per strain |
|---|---|---|---|---|---|---|---|---|---|---|---|
| 5.0 | 0.24 | 0.99 | 0.44 | 0.17 | 0.01 | 0.34 | 0.28 | 0.092 | 0.4 | 0.07 | 0.3 |
| 5.5 | 0.33 | 0.81 | 0.54 | 0.15 | 0.075 | 0.45 | 0.3 | 0.13 | 0.42 | 0.061 | 0.33 |
| 6.0 | 0.38 | 0.87 | 0.46 | 0.1 | 0.18 | 0.31 | 0.31 | 0.094 | 0.42 | 0.11 | 0.32 |
| 6.5 | 0.4 | 0.7 | 0.29 | 0.05 | 0.4 | 0.21 | 0.33 | 0.037 | 0.36 | 0.12 | 0.29 |
| 7.0 | 0.24 | 0.75 | 0.27 | 0.024 | 0.45 | 0.086 | 0.3 | 0.077 | 0.36 | 0.17 | 0.27 |
| 7.5 | 0.26 | 0.72 | 0.25 | 0.2 | 0.43 | 9.1e-3 | 0.31 | 0.17 | 0.25 | 0.28 | 0.29 |
| 8.0 | 0.29 | 0.6 | 0.26 | 0.2 | 1.2 | 0.043 | 0.29 | 0.095 | 0.32 | 0.29 | 0.36 |
| Mean per ROI | 0.31 | 0.78 | 0.36 | 0.13 | 0.4 | 0.21 | 0.3 | 0.1 | 0.36 | 0.16 | |

Table 10: Error of the model's prediction for E' in the adult hippocampus.

| $0.2\sqrt{R\delta}$/R | ML | GCL | hilus | SO3 | SP3 | SR3 | SLM | SR1 | S | Alveus | Mean per strain |
|---|---|---|---|---|---|---|---|---|---|---|---|
| 5.0 | 0.37 | 1.4 | 0.47 | 0.15 | 0.17 | 0.31 | 0.25 | 0.15 | 0.46 | 0.071 | 0.38 |
| 5.5 | 0.42 | 1.2 | 0.48 | 0.096 | 0.15 | 0.36 | 0.23 | 0.19 | 0.42 | 0.047 | 0.35 |
| 6.0 | 0.48 | 1.1 | 0.47 | 0.08 | 0.15 | 0.24 | 0.28 | 0.14 | 0.43 | 0.14 | 0.35 |
| 6.5 | 0.5 | 1.1 | 0.37 | 0.11 | 0.39 | 0.25 | 0.29 | 0.16 | 0.37 | 0.047 | 0.36 |
| 7.0 | 0.36 | 1.0 | 0.38 | 0.046 | 0.27 | 0.17 | 0.3 | 0.062 | 0.38 | 0.14 | 0.31 |
| 7.5 | 0.44 | 0.84 | 0.35 | 0.14 | 0.35 | 0.068 | 0.32 | 6.0e-3 | 0.3 | 0.25 | 0.31 |
| 8.0 | 0.44 | 0.83 | 0.34 | 0.17 | 0.91 | 0.14 | 0.23 | 0.076 | 0.39 | 0.24 | 0.38 |
| Mean per ROI | 0.43 | 1.1 | 0.41 | 0.11 | 0.34 | 0.22 | 0.27 | 0.11 | 0.39 | 0.14 | |

Table 11: Error of the model's prediction for E" in the adult hippocampus.

#### 12 Analysis of E' and E" in juvenile samples of the hippocampus: Affine model

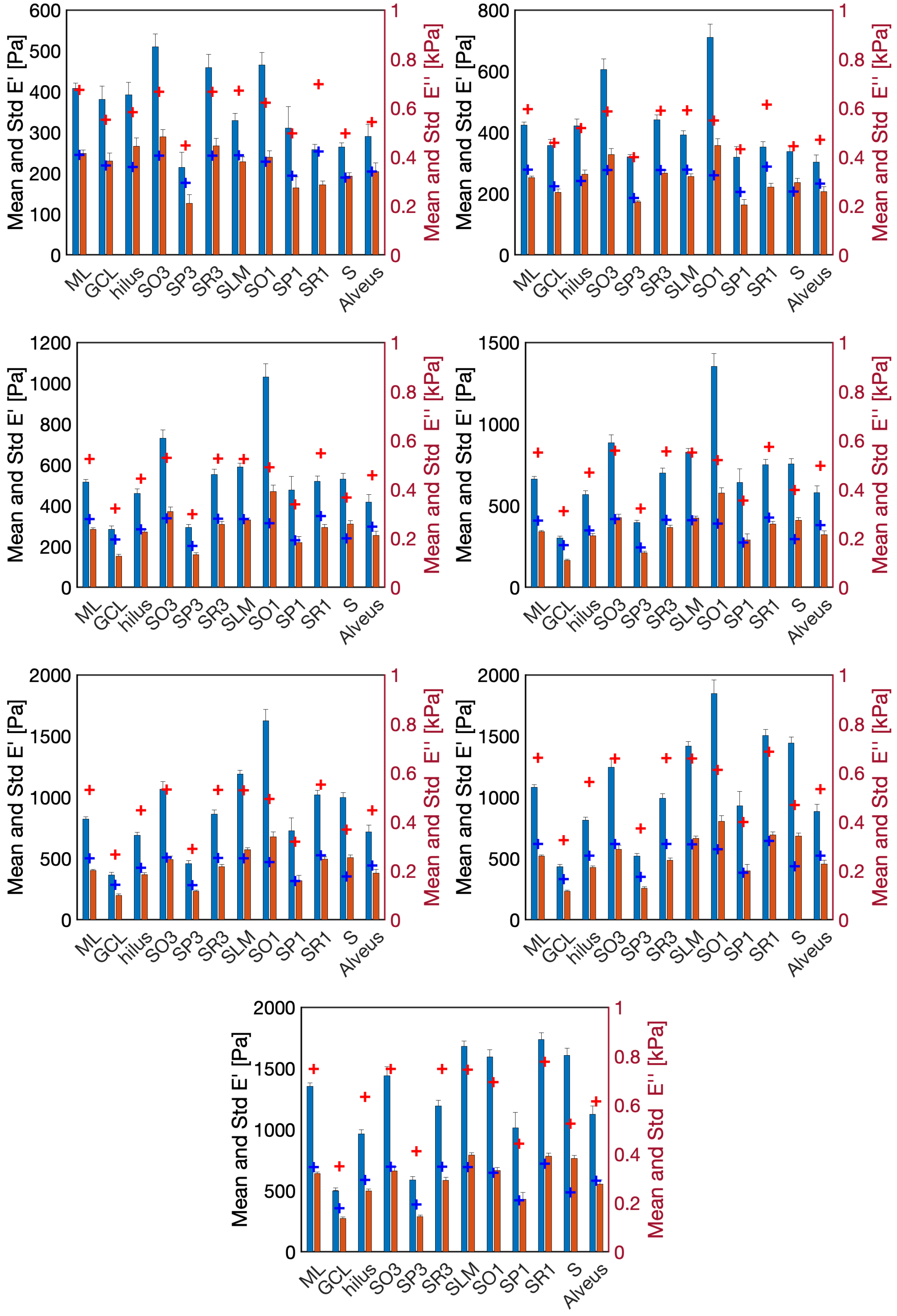


Figure 11: Stiffness values at different ROIs of juvenile mice. Figures show from top to bottom and left to right the results for 7 indentation strain levels: 5, 5.5, 6, 6.5, 7, 7.5 and 8%, respectively. Mean and standard deviation of E' and E" for the different regions of the hippocampus. + marks shows the prediction of the affine mechanical model.

| $0.2\sqrt{R\delta}$/R | ML | GCL | hilus | SO3 | SP3 | SR3 | SLM | SO1 | SP1 | SR1 | S | Alveus | Mean per strain |
|---|---|---|---|---|---|---|---|---|---|---|---|---|---|
| 5.0 | 9.7e-3 | 0.13 | 0.11 | 0.22 | 0.25 | 0.13 | 0.22 | 0.2 | 0.042 | 0.62 | 0.13 | 0.12 | 0.18 |
| 5.5 | 0.12 | 0.022 | 0.017 | 0.23 | 3.6e-3 | 0.067 | 0.21 | 0.38 | 0.083 | 0.4 | 0.052 | 0.24 | 0.15 |
| 6.0 | 0.22 | 0.36 | 0.15 | 0.13 | 0.22 | 0.14 | 0.065 | 0.43 | 0.15 | 0.26 | 0.17 | 0.31 | 0.22 |
| 6.5 | 0.24 | 0.55 | 0.24 | 0.055 | 0.22 | 0.19 | 4.2e-3 | 0.42 | 0.17 | 0.14 | 0.21 | 0.28 | 0.23 |
| 7.0 | 0.29 | 0.45 | 0.3 | 3.7e-3 | 0.26 | 0.23 | 0.11 | 0.39 | 0.12 | 0.083 | 0.26 | 0.24 | 0.23 |
| 7.5 | 0.22 | 0.5 | 0.39 | 0.057 | 0.44 | 0.33 | 0.072 | 0.34 | 0.14 | 0.089 | 0.35 | 0.21 | 0.26 |
| 8.0 | 0.11 | 0.4 | 0.31 | 0.038 | 0.4 | 0.25 | 0.11 | 0.13 | 0.13 | 0.1 | 0.35 | 0.093 | 0.2 |
| Mean per ROI | 0.17 | 0.34 | 0.22 | 0.1 | 0.26 | 0.19 | 0.11 | 0.33 | 0.12 | 0.24 | 0.22 | 0.21 | |

Table 12: Error of the model's prediction for E' in the juvenile hippocampus.

| $0.2\sqrt{R\delta}$/R | ML | GCL | hilus | SO3 | SP3 | SR3 | SLM | SO1 | SP1 | SR1 | S | Alveus | Mean per strain |
|---|---|---|---|---|---|---|---|---|---|---|---|---|---|
| 5.0 | 0.017 | 0.049 | 0.19 | 0.16 | 0.4 | 0.088 | 0.07 | 0.043 | 0.18 | 0.47 | 0.019 | 2.1e-3 | 0.14 |
| 5.5 | 0.11 | 0.099 | 0.082 | 0.15 | 0.071 | 0.038 | 0.085 | 0.28 | 0.26 | 0.3 | 0.13 | 0.12 | 0.14 |
| 6.0 | 0.17 | 0.53 | 0.047 | 0.093 | 0.26 | 0.088 | 0.012 | 0.33 | 0.05 | 0.19 | 0.23 | 0.17 | 0.18 |
| 6.5 | 0.2 | 0.57 | 0.1 | 0.019 | 0.15 | 0.13 | 0.033 | 0.33 | 0.056 | 0.11 | 0.28 | 0.19 | 0.18 |
| 7.0 | 0.25 | 0.42 | 0.15 | 0.035 | 0.21 | 0.16 | 0.12 | 0.31 | 5.3e-3 | 0.062 | 0.31 | 0.16 | 0.18 |
| 7.5 | 0.19 | 0.43 | 0.23 | 0.074 | 0.36 | 0.27 | 0.074 | 0.28 | 0.04 | 0.074 | 0.36 | 0.15 | 0.21 |
| 8.0 | 0.084 | 0.31 | 0.18 | 0.05 | 0.34 | 0.19 | 0.13 | 0.029 | 0.021 | 0.076 | 0.36 | 0.052 | 0.15 |
| Mean per ROI | 0.15 | 0.34 | 0.14 | 0.083 | 0.26 | 0.14 | 0.074 | 0.23 | 0.088 | 0.18 | 0.24 | 0.12 | |

Table 13: Error of the model's prediction for E" in the juvenile hippocampus.

#### 13 Analysis of E’ and E” in juvenile samples of the cerebellum: Affine model

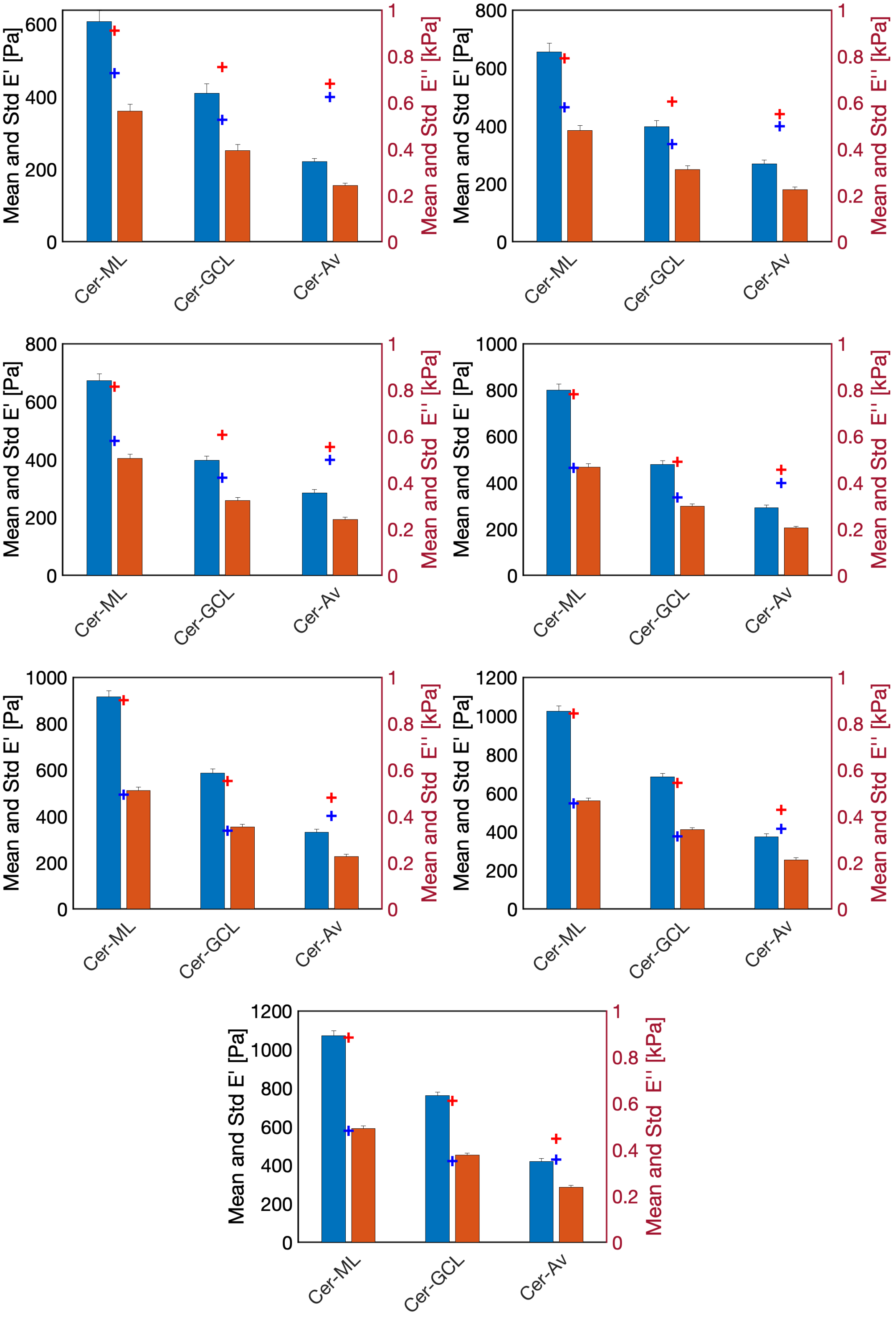


Figure 12: Stiffness values at different ROIs of juvenile mice. Figures show from top to bottom and left to right the results for 7 indentation strain levels: 5, 5.5, 6, 6.5, 7, 7.5 and 8%, respectively. Mean and standard deviation of E’ and E” for the different regions of the cerebellum. + marks shows the prediction of the affine mechanical model.

| $0.2\sqrt{R\delta}$/R | ML | GCL | Alveus | Mean per strain |
|---|---|---|---|---|
| 5.0 | 0.041 | 0.18 | 0.97 | 0.4 |
| 5.5 | 0.033 | 0.22 | 0.64 | 0.3 |
| 6.0 | 0.03 | 0.22 | 0.55 | 0.27 |
| 6.5 | 0.022 | 0.025 | 0.56 | 0.2 |
| 7.0 | 0.015 | 0.058 | 0.45 | 0.17 |
| 7.5 | 0.013 | 0.047 | 0.37 | 0.14 |
| 8.0 | 0.01 | 0.035 | 0.28 | 0.11 |
| Mean per ROI | 0.023 | 0.11 | 0.55 | |

Table 14: Error of the model's prediction for E' in the juvenile cerebellum.

| $0.2\sqrt{R\delta}$/R | ML | GCL | Alveus | Mean per strain |
|---|---|---|---|---|
| 5.0 | 0.29 | 0.34 | 1.6 | 0.73 |
| 5.5 | 0.21 | 0.35 | 1.2 | 0.59 |
| 6.0 | 0.15 | 0.3 | 1.1 | 0.51 |
| 6.5 | 4.9e-3 | 0.13 | 0.95 | 0.36 |
| 7.0 | 0.033 | 0.048 | 0.77 | 0.28 |
| 7.5 | 0.026 | 0.089 | 0.63 | 0.25 |
| 8.0 | 0.022 | 0.072 | 0.5 | 0.2 |
| Mean per ROI | 0.11 | 0.19 | 0.96 | |

Table 15: Error of the model's prediction for E" in the juvenile cerebellum.

### 14 Changes in E' and E" along indentation strain values in all samples: Affine model

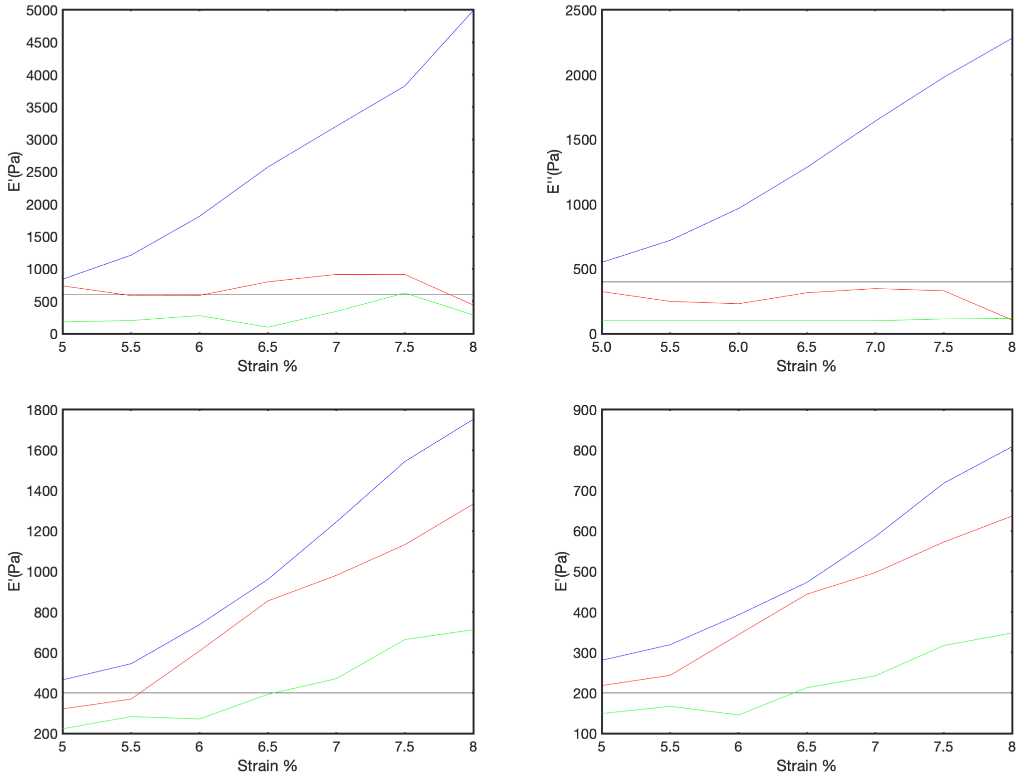


Figure 13: Evolution of E' (left) and E" (right) for adult (top) and juvenile mice (bottom) as a function of the indentation strains, 5.5, 6, 6.5, 7, 7.5 and 8%, of the affine model. Black, red, green and blue curves for cell bodies, myelin, PNN and NIM, respectively.

### 15 Comparison adult and juvenile stiffness prediction

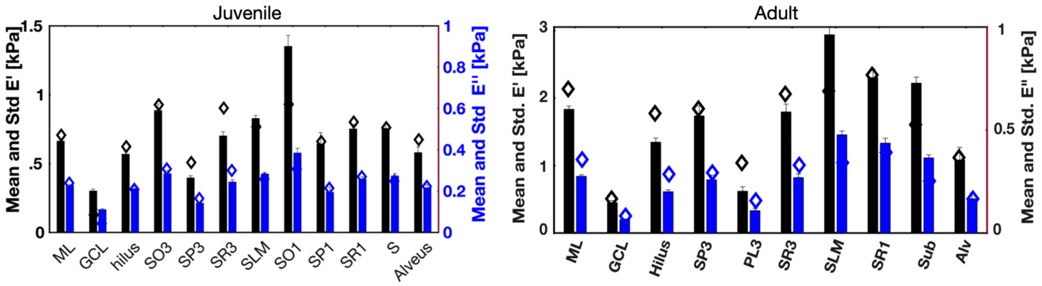


Figure 14: Comparison between juvenile (left) and adult (right) stiffness values and model predictions.

#### 16 Description of ROIs

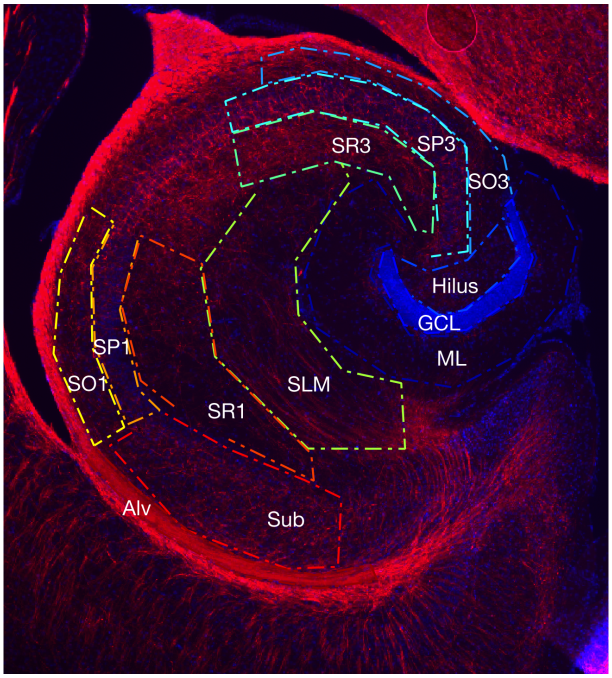


Figure 15: The 12 regions of interest delimited by dash-point lines in a stained images of the hippocampus. Abbreviations used for regions are: Alv - alveus, Sub - subiculum, SLM - stratum lacunosum moleculare, SR - stratum radiatum, SP - stratum pyramidale, SO - stratum oriens, ML - molecular layer, GCL - granule cell layer.
